# Contrasting evolutionary trajectories of nitrate assimilation across *Brettanomyces bruxellensis* lineages

**DOI:** 10.64898/2026.08.31.748220

**Authors:** Audrey Vigna, Jules Harrouard, Cecile Miot-Sertier, Louise Michelizza, Victor Loegler, Philippe Marullo, Anne Friedrich, Joseph Schacherer, Emilien Peltier, Warren Albertin

## Abstract

*Brettanomyces bruxellensis* is a yeast species associated with diverse fermentation environments and characterized by extensive genetic diversity, including diploid, autotriploid, and allotriploid lineages resulting from independent hybridization events. These lineages are associated with distinct ecological niches and provide a framework for studying metabolic trait evolution in complex genomes. Nitrate assimilation is a relatively uncommon trait among yeasts and has been reported in *B. bruxellensis*, but its distribution and evolutionary history within the species remain poorly understood. Here, we combined phenotypic characterization of 151 strains with genomic analyses of 946 whole-genome sequences to investigate nitrate assimilation. Growth assays revealed that nitrate assimilation is widespread but unevenly distributed across genetic lineages, with some populations largely retaining the trait whereas others have frequently lost it. Genomic analyses identified extensive variation affecting the nitrate assimilation gene cluster composed of *YNR1*, *YNI1*, and *YNT1*. Nitrate assimilation was strongly associated with both gene copy number and predicted gene functionality, with nitrate-assimilating strains generally carrying more functional copies of the cluster. Leveraging the complex genomic architecture of the species, we independently analyzed primary and acquired genomes in allotriploid lineages and uncovered contrasting evolutionary trajectories following hybridization. While nitrate assimilation genes were generally maintained in primary genomes, acquired genomes showed a higher prevalence of gene loss and predicted loss-of-function variants, revealing asymmetric dynamics between subgenomes. Altogether, our results suggest that nitrate assimilation represents an ancestral trait that has been differentially maintained across *B. bruxellensis* lineages through a combination of copy number variation, gene degeneration, and genome-specific evolutionary dynamics. These findings provide new insights into how genome architecture and polyploid evolution shape the maintenance and loss of metabolic traits in an industrially relevant yeast species.

**Author Summary:** Microorganisms adapt to new environments by gaining, modifying, or losing biological functions over evolutionary time. Understanding how these processes occur remains a central question in evolutionary biology, particularly in species with complex genomes. The yeast *Brettanomyces bruxellensis* provides an interesting model because it is associated with diverse fermentation environments and genetically distinct lineages with different evolutionary histories. We focused on nitrate assimilation, a relatively uncommon ability among yeasts that enables the use of nitrate as a nitrogen source. By combining growth experiments with large-scale genome analyses, we investigated how this trait is distributed across the species and how it evolved. We found that nitrate assimilation is likely an ancestral trait, but that it has been maintained or lost depending on the lineage. These differences are associated with variation in the number and integrity of genes involved in nitrate assimilation. *B. bruxellensis* also contains hybrid lineages carrying both primary and acquired genomes. By analyzing these genomes separately, we found that nitrate assimilation genes follow different evolutionary trajectories, with acquired genomes showing a greater tendency toward gene loss. Our results illustrate how hybridization and genome architecture can influence the maintenance or loss of biological functions within a species.

## Introduction

The budding yeast *Brettanomyces bruxellensis* is recognized as a major contaminant in industrial fermentation processes, particularly in beverage fermentations such as wine and beer, as well as in bioethanol production (1). In the wine sector, it is associated with the production of volatile phenols responsible for organoleptic deviations commonly referred to as “taste of Brett”, characterized by sweaty, leathery, or horse-like notes to the product (2). In bioethanol production processes, its presence is mainly correlated with reduced ethanol yields, resulting in significant economic losses (3). In brewing, its impact appears more variable, ranging from spoilage of conventional beers to the sought-after contribution of aromatic compounds in certain traditional styles, such as sour beers (4–6). Beyond its classification as a spoilage organism, some studies have also explored the potential use of *B. bruxellensis* in industrial fermentation processes (7).

In addition to its biotechnological impact, *B. bruxellensis* constitutes a particularly valuable model organism due to its remarkable genetic complexity (8–11). Previous studies have revealed a clear population structure comprising seven genetically distinct clusters, notably differing in ploidy. Three clusters are strictly diploid and are considered representative of the ancestral state of the species. However, more than half of the described isolates are triploid, arising from independent intra-or interspecific hybridization events (8,9,11,12). Three of these clusters result from allotriploidization events involving *B. bruxellensis* and three yet-uncharacterized species exhibiting more than 3% genetic divergence. These events led to the acquisition of an additional haploid genome, distinct from the primary diploid genome, thereby increasing genetic diversity and adaptive potential. Another cluster originated from an autotriploidization event resulting from hybridization between two *B. bruxellensis* lineages (10,11).

The genetic structuring of these lineages is closely associated with the ecological niches and geographical contexts from which the strains were isolated, suggesting processes of local adaptation (9–11). In this context, polyploidization, considered one of the most extreme mutational events in nature, could play a central role in the evolution of the species by promoting stress tolerance and adaptation to fluctuating environments (13). This genetic diversity is further reinforced by copy number variations (CNVs) and loss of heterozygosity (LOH) events, which are likely to shape key physiological traits (11).

The spoilage ability of *B. bruxellensis* largely relies on its scavenger-type metabolism, enabling growth in nutrient-poor environments, typically after the main alcoholic fermentation stages carried out notably by *Saccharomyces* cerevisiae (4,14,15). The species is particularly adapted to conditions of high ethanol concentration, low pH, and the presence of sulfites with more than 40% of isolates displaying resistance (9,16–19). These traits are supported by a remarkable ability to exploit a wide range of carbon and nitrogen sources, depending on oxygen availability and cofactor supply (7,8,15,16,20–23).

Among nitrogen sources, nitrate assimilation has been reported in *B. bruxellensis*, although this trait appears heterogeneous across the species and remains poorly documented (16,20,21,24–26). Nitrate-assimilating yeasts are rare in nature, as this metabolism is generally considered energetically costly and prone to frequent evolutionary loss (27). Nitrate assimilation involves three successive steps: nitrate transport across the membrane, followed by two enzymatic reduction steps leading to the production of ammonium, as described in *Hansenula polymorpha*. These processes are mediated by the NIT gene cluster, composed of three adjacent genes, *YNT1*, *YNI1*, and *YNR1* (28). In addition, two transcriptional cofactors, *YNA1* and *YNA2*, have been identified in nitrate-assimilating yeasts (29). In *B. bruxellensis*, orthologs of the nitrate assimilation genes (*YNT1, YNI1, YNR1*) have been identified (30,31).

In this study, we investigate the ability of *B. bruxellensis* to assimilate nitrate as a sole nitrogen source under aerobic conditions, characterize the distribution of this trait across the species, and identify its underlying genetic determinants. By integrating whole-genome sequencing data from a collection of more than 1,000 isolates with an extensive phenotypic screening of 151 strains, we examine the relationship between nitrate assimilation and the complex population and genomic structure of the species. In particular, we investigate how variation in ploidy, genomic architecture, and the distinct evolutionary trajectories of subgenomes have shaped this trait across the diverse lineages and ecological niches associated with *B. bruxellensis*.

## Results

### 1. Nitrate assimilation varies extensively across *B. bruxellensis* strains

To investigate the phenotypic variation in nitrate assimilation in *B. bruxellensis*, we analyzed a panel of 151 strains selected to represent the genetic diversity and population structure of the species across the different genetic clusters (Fig. 10; see Materials and Methods). The ability of each strain to assimilate nitrate was assessed using growth assay performed in three synthetic media differing in their nitrogen source: YPD as a rich control medium, and two minimal media containing either ammonium (NH₄⁺) or nitrate (NO₃⁻) as the sole nitrogen source. These media are hereafter referred to as YNB+A and YNB+N, respectively. Growth was monitored in 96-well microplates under aerobic conditions with three biological replicates per strain and condition, resulting in more than 1,500 growth curves. Analysis of the growth curves revealed three distinct growth behaviors (Fig 1A): strain able to grow in all tested conditions (YPD, YNB+A and YNB+N), strain able to grow on YPD and YNB+A but unable to grow on nitrate, and a last group of strains that grew exclusively on YPD. Growth was defined as a final optical density exceeding 0.5 OD. These phenotypes were classified as NIT⁺ (growth on all media, including YNB+N), NIT⁻ (no growth on YNB+N but growth on YPD and YNB+A), and AMMO⁻ (no growth on YNB+A, despite growth on YPD and YNB+N), respectively. Representative growth profiles for each of these three phenotypic categories are shown in Figure 1A. Overall, across the 151 phenotyped strains, nitrate-consuming (NIT⁺), nitrate-non-consuming (NIT⁻), and ammonium-non-consuming (AMMO⁻) phenotypes accounted for 65% (n = 98), 19% (n = 29), and 16% (n = 24) of the strains, respectively (Fig. 1B).

**Fig. 1.**
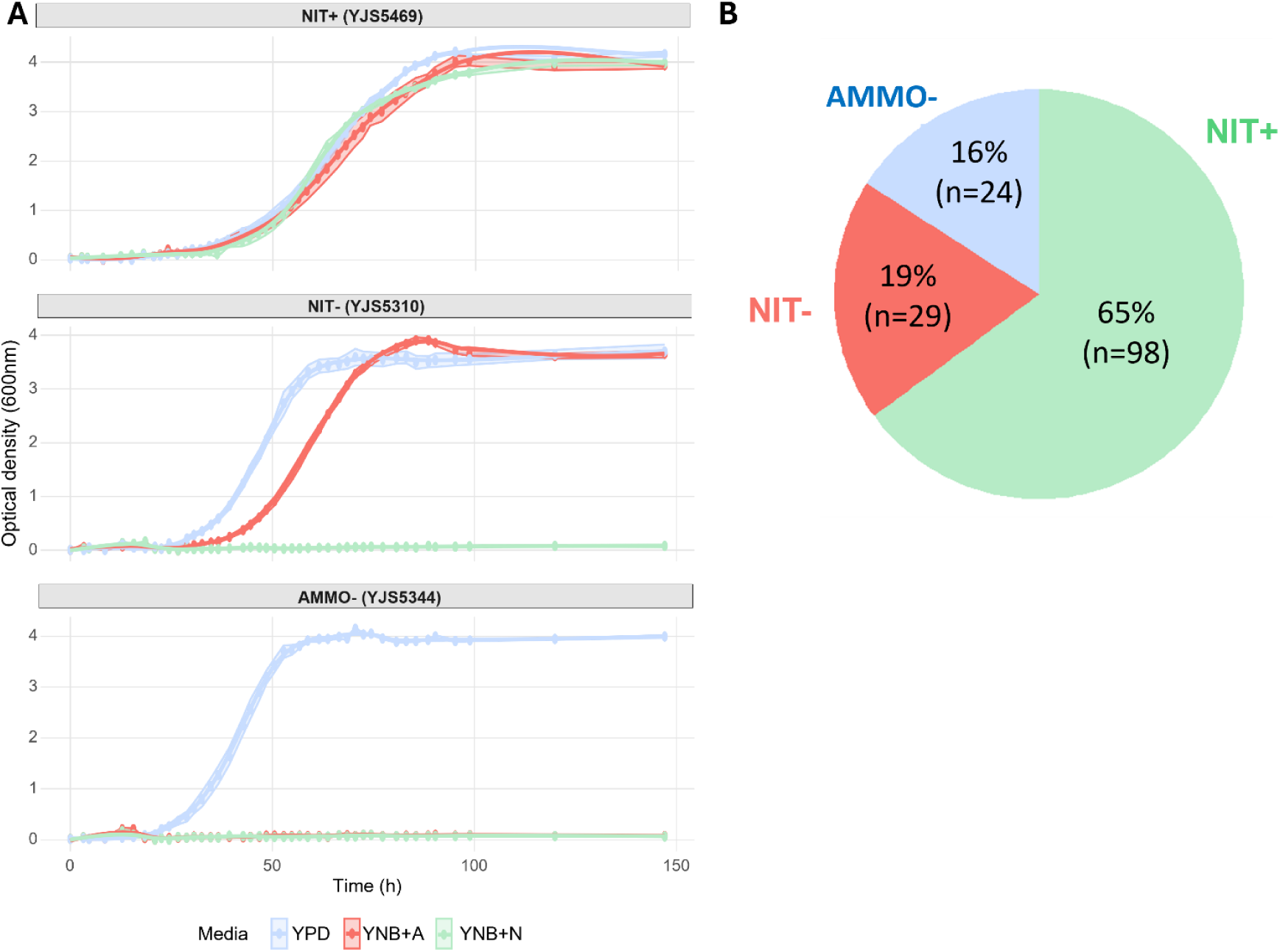
Growth phenotypes and quantitative growth parameters of *B. bruxellensis* on different nitrogen sources. **(A)** Representative growth curves illustrating the three phenotypic classes identified based on nitrogen source utilization. Optical density (OD₆₀₀) was monitored over 150 h under aerobic conditions in three media: YPD, YNB+A (minimal medium with ammonium as the sole nitrogen source), and YNB+N (minimal medium with nitrate as the sole nitrogen source). One representative strain is shown for each phenotype: NIT⁺ (growth on all three media), NIT⁻ (growth on YPD and YNB+A but not on YNB+N), and AMMO⁻ (growth on YPD and YNB+N but not on YNB+A). Solid lines represent the mean of three biological replicates, and shaded areas indicate the associated variability. **(B)** Pie chart of the relative proportions of the three nitrogen phenotypes (NIT⁺, NIT⁻, and AMMO⁻), with percentages and corresponding strain counts indicated.

These results reveal marked phenotypic variations in nitrogen source utilization across *B. bruxellensis*. We therefore next examined how these phenotypes are distributed across the genetic clusters of the species.

### 2. Nitrate assimilation varies across the genetic clusters of *B. bruxellensis*

The distribution of nitrogen-related phenotypes across the genetic clusters was first examined using a contingency table analysis (Fig. 2A). In Fig. 2A, filled circles represent observed counts, while open circles indicate expected counts under the hypothesis of homogeneous distribution amongst population. A chi-squared test of independence revealed a highly significant association between phenotype and genetic structure (χ² test, p= 9.583e-12), indicating that nitrate assimilation phenotypes are not randomly distributed across the species. Cluster-specific chi-square goodness-of-fit tests were used to assess deviations from the global phenotype distribution, with p-values corrected for multiple testing (Benjamini–Hochberg). After correction, the Non-admixed D2 and Admixed D1/D2 clusters showed the strongest deviations from the rest of the species, consistent with their reduced frequency of NIT⁺ strains. Nitrate-consuming strains (NIT⁺) are present in all genetic clusters, but with an uneven distribution. Some clusters show a high proportion of nitrate-assimilating strains, such as diploid D1 with 57% of NIT⁺ strains and Admixed D2 with 100% of the tested strains classified as NIT⁺, as well as the three allotriploid groups A1, A2 and A3, for which the proportion of NIT⁺ strains exceeds 70%. In contrast, other clusters show a markedly lower prevalence of this phenotype, notably the autotriploid Admixed D1/D2 cluster, in which more than 65% of strains lack this phenotype. The inability to grow on ammonium phenotype (AMMO⁻) was restricted to specific clusters, such as diploid D1, where 20% of the tested strains were unable to consume ammonium, diploid Non-admixed D2, where 70% of the strains were classified as AMMO⁻, and the allotriploid A2 cluster, in which 25% of strains exhibited this phenotype (Fig. 2A).

**Fig. 2.**
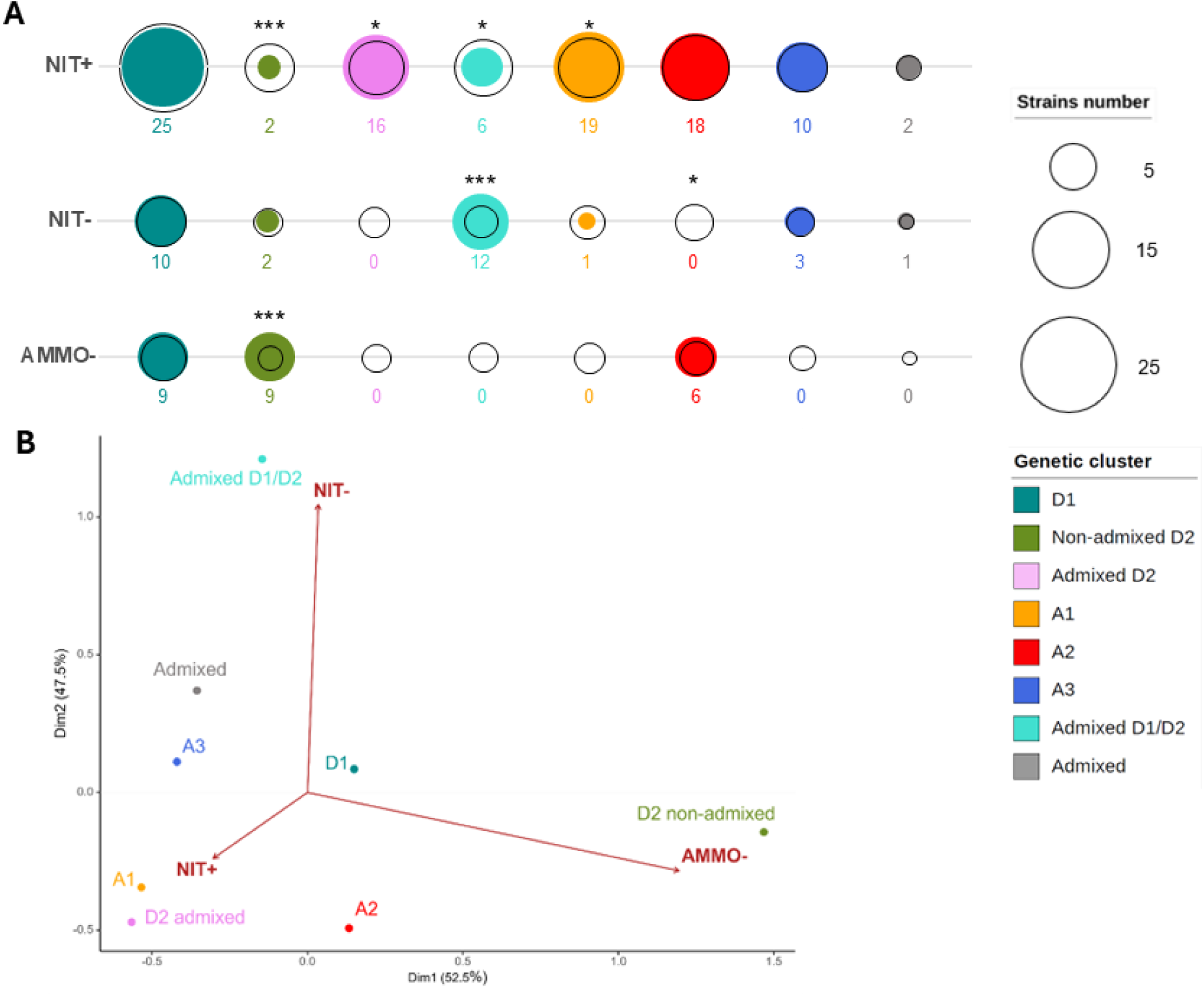
Distribution of nitrogen-related phenotypes across *B. bruxellensis* (n = 151 strains). **(A)** Distribution of nitrogen-related phenotypes (AMMO−, NIT−, NIT+) across genetic clusters. Filled circles represent observed counts, whereas open circles indicate expected counts under the null hypothesis of independence. Circle size is proportional to the number of strains exhibiting each phenotype within a given cluster, and numbers correspond to the observed strain counts. Colors indicate genetic clusters. A global chi-square test of independence indicates a significant association between phenotype and genetic clusters (*χ²*, *p* = 9.583e-12). Asterisks denote phenotype × cluster combinations showing significant deviations from the expected distribution, based on standardized residuals from the chi-square test with Benjamini–Hochberg correction (*p*adj < 0.05: *, < 0.01: **, < 0.001: ***). **(B)** Biplot of correspondence analysis (CA) summarizing associations between genetic clusters and nitrogen-related phenotypes. Genetic clusters are represented as colored points, while phenotypes are shown as vectors in the reduced ordination space.

Correspondence analysis (CA) was then applied to the phenotype × genetic cluster contingency table to visualize associations between genetic clusters and nitrogen-related phenotypes (Fig. 2B). Using symmetric scaling, CA revealed a strong structuring of genetic clusters according to nitrogen-related phenotypes. The first dimension (Dim1, 52.5% of total inertia) separates clusters associated with ammonium non-utilization (AMMO−) from those associated with nitrate assimilation. The second dimension (Dim2, 47.5% of inertia) further discriminates clusters according to the presence or absence of nitrate assimilation. The NIT⁺ phenotype was predominantly associated with allotriploid clusters as well as D1 and Admixed D2 clusters, and was positioned opposite the autotriploid Admixed D1/D2 cluster along the second dimension, in which nitrate assimilation was rare. The AMMO⁻ phenotype was strongly associated with the Non-admixed D2 cluster along the first dimension. Both the contingency-based analyses and correspondence analysis indicate that nitrogen utilization phenotypes are strongly structured by the genetic structure of *B. bruxellensis*.

### 3. Predicted competitive fitness reflects nitrate assimilation capacity and population structure

To assess whether nitrate assimilation is associated with differences in fitness beyond nitrate utilization itself, we compared quantitative growth parameters between nitrate-assimilating (NIT⁺) and non-assimilating (NIT⁻) strains in media not containing nitrate (YNB+A and YPD). Growth in YNB+N was analyzed separately as a validation of the nitrate assimilation phenotype. The maximal optical density, maximal growth rate (rₘₐₓ), and lag phase were used for these comparisons (Fig. 3). AMMO⁻ strains were excluded from this analysis, as this phenotype reflects ammonium auxotrophy rather than the absence of nitrate assimilation and could therefore confound the interpretation. In both YNB+A and YPD, no significant difference in maximal optical density was observed between NIT⁺ and NIT⁻ strains, with mean values ranging from approximately 3.7 to 3.9 regardless of phenotype. Similarly, maximal growth rates were comparable between phenotypes in YNB+A (mean rₘₐₓ ≈ 0.12 h⁻¹). In YPD, NIT⁺ strains exhibited a slightly higher maximal growth rate than NIT⁻ strains (mean rₘₐₓ = 0.14 h⁻¹ vs 0.13 h⁻¹), although the difference remained small. Lag phase duration did not differ significantly between NIT⁺ and NIT⁻ strains in either medium. As expected, NIT⁻ strains did not grow in YNB+N, whereas NIT⁺ strains reached maximal optical density values comparable to those observed in other media (≈ 3.7), with a mean maximal growth rate of approximately 0.11 h⁻¹.

**Fig. 3.**
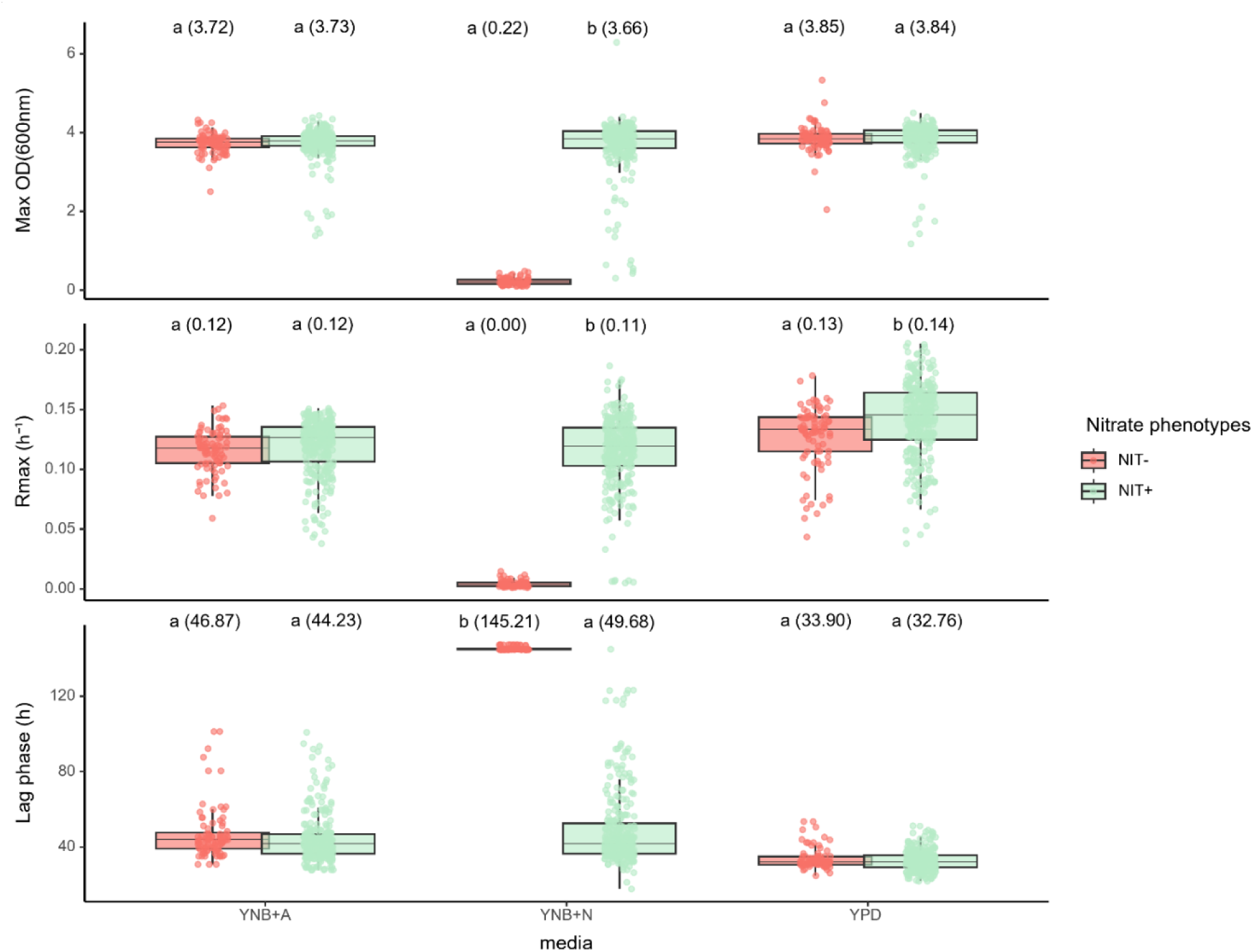
Comparison of quantitative growth parameters between nitrate-assimilating and non-assimilating strains across nitrogen sources. Boxplots show the distribution of maximal optical density, maximal growth rate (rₘₐₓ), and lag phase duration for nitrate-assimilating (NIT⁺) and non-assimilating (NIT⁻) strains across YNB+A, YNB+N, and YPD media. Growth parameters were estimated from generalized additive model (GAM) fits of growth curves. AMMO⁻ strains were excluded from the analysis. Points represent strain-specific values. For each parameter and medium, statistical comparisons between NIT⁺ and NIT⁻ strains were performed using linear mixed-effects models including phenotype, medium, and their interaction as fixed effects, and strain identity as a random effect. Different letters indicate statistically significant differences between groups within a given medium based on post hoc pairwise comparisons (p < 0.05).

To determine whether slight differences in growth performance observed in rich medium translate into individual differences in inferred competition, we analyzed the outcomes of *in silico* pairwise growth simulations based on parameters estimated from monocultures. For each pairwise simulation, competition fitness was quantified as the level of colonization of the medium, expressed as a percentage (eg 50% means both strains showed similar growth, 0% and 100% means one strain was fully outcompeted by the other). Strain-level performance therefore corresponds to the mean colonization obtained across all pairwise simulations within a given medium (Fig. 4). Given that nitrate assimilation phenotypes are strongly structured by the genetic background of the species (Fig. 3), we further examined these competitive outcomes at the level of genetic clusters.

**Fig. 4.**
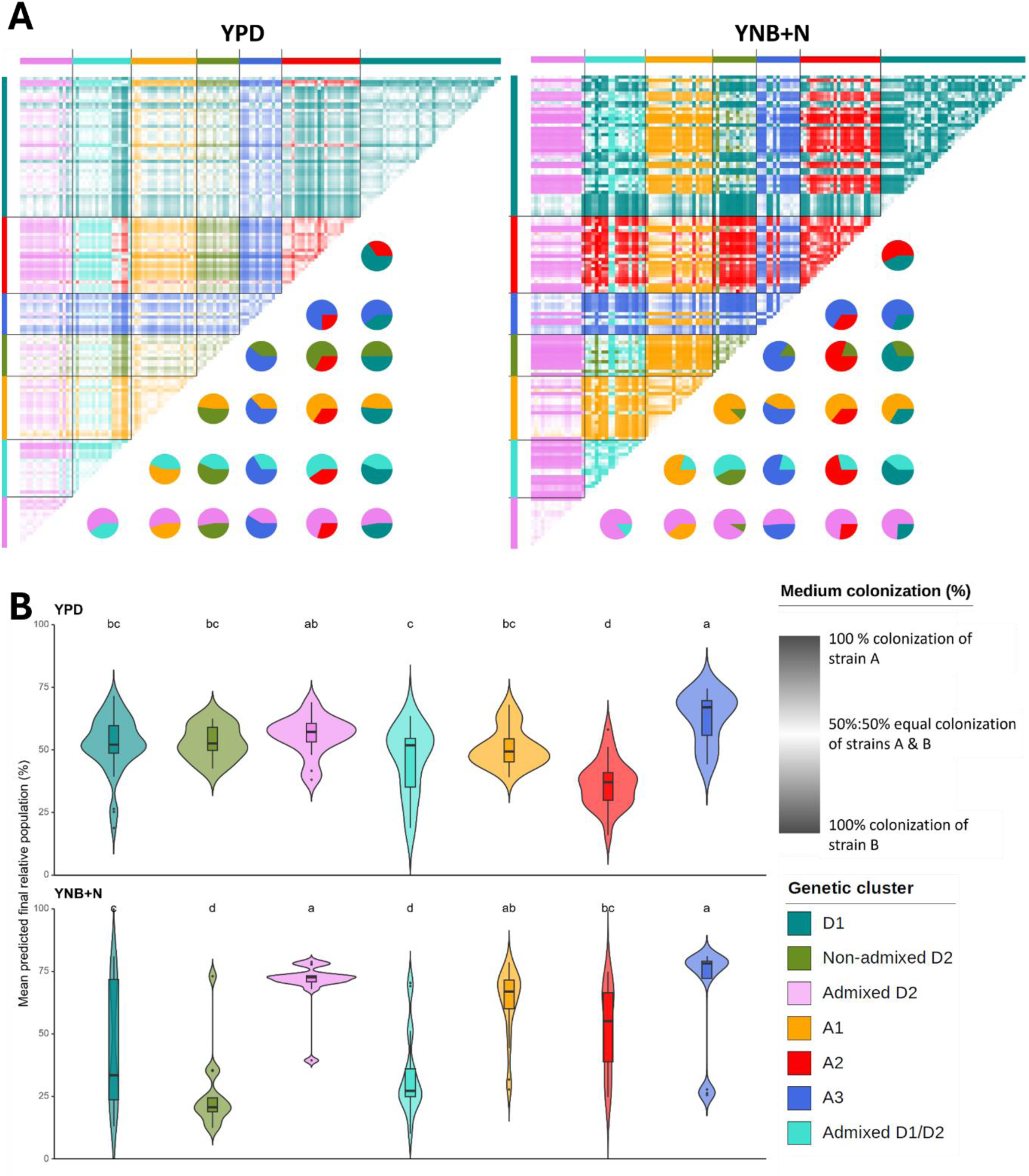
*In silico* pairwise competitions across nitrogen sources and genetic clusters (n=151 strains). **(A)** Pairwise simulation matrices showing predicted final relative population fractions between all pairs of strains grown in YNB+N and YPD media. Simulations were performed for the full set of 151 strains and were based exclusively on growth parameters calculated from monocultures. Each cell represents the predicted final relative population fraction between a pair of strains at the end of the simulation window. Cell color intensity reflects the magnitude of asymmetry in their predicted competitions: pale colors indicate near-equal outcomes (∼50/50), whereas increasingly saturated colors indicate stronger deviations equal colonization. Color identity indicates which of the two strains reached the higher final relative population level. Associated pie charts summarize, for each strain, the distribution of competitions against strains from different genetic clusters. **(B)** Distribution of strain-level predicted relative population values across genetic clusters in YNB+N and YPD. For each strain, mean predicted competition performance was calculated as the average final relative population fraction obtained across all pairwise simulations within the same medium. Violin plots show the distribution of these values within each genetic cluster. Different letters denote statistically significant differences among genetic clusters within a given medium based on Kruskal–Wallis tests followed by post hoc multiple comparisons (p < 0.05).

In the rich medium YPD, pairwise competition matrices (Fig. 4A) displayed values broadly centered around 50%, with no clear fitness pattern emerging between strains. Cluster-level pie charts likewise showed relatively balanced outcomes overall, although subtle deviations were observed: strains from the allotriploid A3 cluster tended to outcompete strains from other clusters more frequently, whereas A2 strains appeared to contribute a lower predicted relative population in pairwise simulations. Violin plot analyses (Fig. 4B) further quantified these patterns. Most genetic clusters exhibited median predicted relative population values close to 50% in YPD. Consistently, the allotriploid A2 cluster showed lower predicted competition ability against most other clusters (median ≈ 34%), whereas A3 strains displayed higher predicted relative population values (median ≈ 68%).

In the minimal medium containing nitrate as the sole nitrogen source (YNB+N), clear cluster-level differences in predicted outcomes emerged. The admixed D2 and A3 genetic clusters reached markedly higher median predicted relative population values (71% and 73%, respectively). The A1 cluster also exhibited an elevated median (≈ 71%), although with greater variability. The A2 cluster showed intermediate and more variable outcomes in YNB+N (median ≈ 53%). The D1 cluster also displayed variable outcomes, with a median of 42%, consistent with the intermediate proportion of NIT+ strains within this cluster (25/44). In contrast, the autotriploid Admixed D1/D2 and the Non-admixed D2 clusters exhibited significantly reduced predicted competitive outcomes in YNB+N, with median values close to 25% and 20%, respectively, in agreement with their low frequency of NIT+ strains.

### 4. Structure and localization of the nitrate assimilation genes in primary and acquired genomes

To investigate the genetic basis of nitrate assimilation, we examined the genomic organization, chromosomal localization, and presence of the nitrate assimilation genes in primary (PG) and acquired (AG) genomes. The putative orthologs of the three nitrate assimilation genes *YNR1*, *YNI1* and *YNT1* in *H. polymorpha* are located on chromosome IV in *B. bruxellensis* reference diploid genome. This chromosome has a total length of 1,751,256 bp (1.75 Mb) and is composed by 733 annotated genes. The three genes involved in nitrate assimilation are organized as a cluster of adjacent genes showing conserved organization and gene order across primary and acquired genomes (Fig.5A). This cluster is located near the left end of chromosome IV in *B. bruxellensis* reference genome. The genomic coordinates and gene length are summarized in Figure 5B. Altogether, the cluster extends from position 17,284 to 26,897 bp, corresponding to a total length of 9.61 kb, including intergenic regions. The nitrate assimilation gene cluster is located ∼17.3 kb from the left end of chromosome IV, placing it within the subtelomeric region. Regarding the presence of these three genes in acquired genomes haploid assemblies (10), comparative analyses revealed that the nitrate assimilation gene cluster is present in the acquired genomes A1 (formerly Beer) and A3 (formerly Teq/EtOH) with a BLAST query coverage of 0.99 for all three genes, but absent from the acquired genome A2 (formerly Wine 1). In A1 and A3 clusters, the three genes were identified based on sequence similarity and annotation transfer, and are organized as a compact gene cluster with conserved gene order relative to the primary genome (Fig. 5B). Analyses of the local genomic context revealed that, in the primary genome as well as in acquired genomes A1 and A3, the nitrate assimilation gene cluster is consistently flanked by the genes *LAC4* and *IMA1*. In contrast, in the acquired genome A2, the three nitrate assimilation genes as well as their two flanking genes (*LAC4* and *IMA1*) are absent. Across all genes located on chromosome IV of the primary genome, the acquired genome A1 covers 97% of primary-genome genes, A3 covers 92%, whereas A2 covers only 85% of primary-genome genes on chromosome IV.

**Fig. 5.**
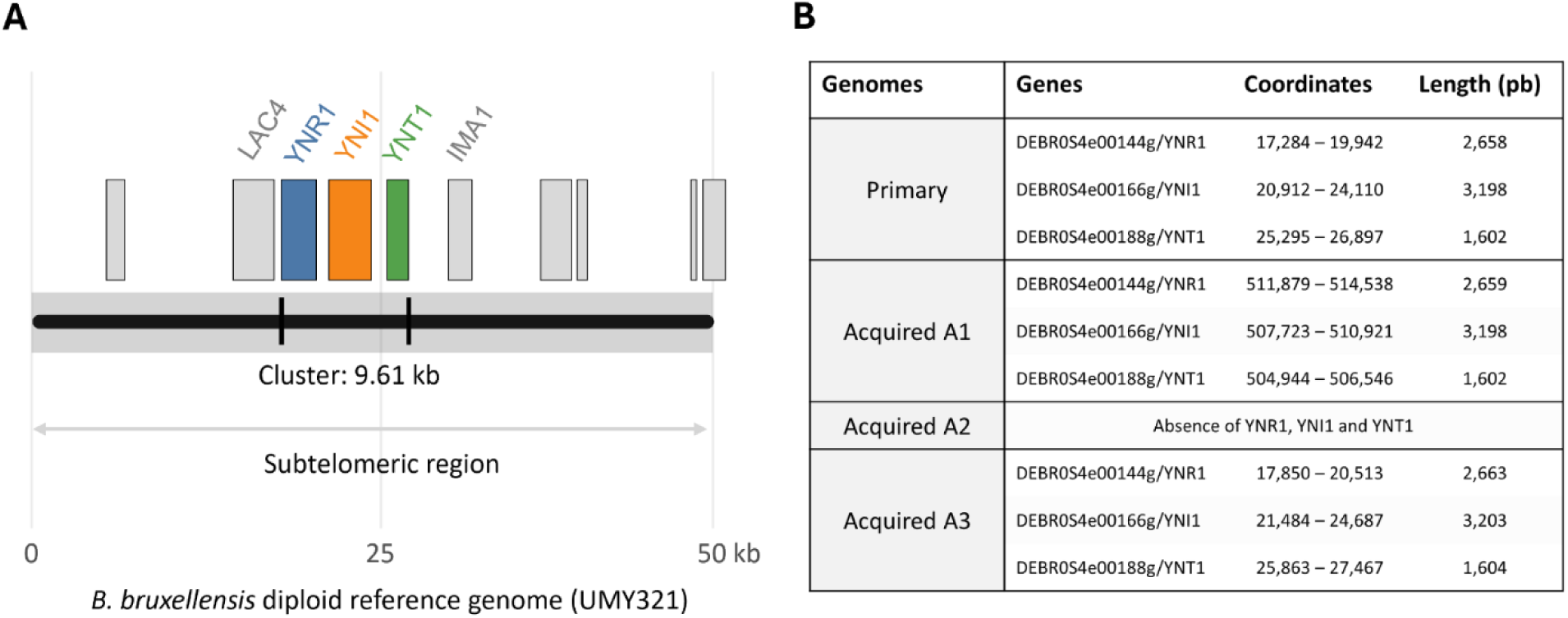
Genomic organization and localization of the nitrate assimilation gene cluster in *B. bruxellensis*. **(A)** Schematic representation of the subtelomeric region of chromosome IV in the *B. bruxellensis* diploid reference genome (UMY321, Fournier et al., 2017). The nitrate assimilation gene cluster, composed of the three adjacent genes *YNR1, YNI1 and YNT1*, is located near the left chromosomal extremity and spans 9.61 kb. The cluster is flanked by the neighboring genes *LAC4* and *IMA1*, which are conserved across genomes in which the cluster is present. Grey boxes represent other gene models located in this subtelomeric region. The shaded region indicates the subtelomeric region (0-50 kb from the chromosome end). Gene positions and distances are shown schematically to illustrate relative organization and local genomic context; the representation is not drawn to scale and may vary among isolates due to structural variation. **(B)** Genomic coordinates and lengths of nitrate assimilation genes in the primary diploid genome and in the acquired genomes A1, A2 and A3. The complete gene cluster is present in the primary genome and in acquired genomes A1 and A3, with conserved gene order and compact organization, whereas all three genes are absent from acquired genome A2. The coordinates of the genes are not comparable between the primary and acquired genomes.

### 5. The nitrate assimilation gene cluster exhibits extensive copy number variation across *B. bruxellensis*

We hypothesized that variation in nitrate assimilation phenotypes could be associated with copy number variation affecting the nitrate assimilation gene cluster. CNV and subsequent genomic analyses were performed on the complete set of 946 sequenced isolates in order to assess genome-wide patterns of variation beyond the 151 phenotyped strains.

To investigate CNV affecting nitrate assimilation genes and to compare it with chromosome-wide copy-number patterns, copy number analyses were performed separately for primary and acquired genomes in allotriploid isolates. We computed for each isolate the mean copy number per gene across the three nitrate assimilation genes (*YNR1*, *YNI1* and *YNT1*), used as a proxy for cluster-level gene dosage, as these genes are physically clustered and shared functional role, and plotted on the vertical axis in Fig. 6A. In parallel, we calculated for each isolate the mean copy number per gene across all genes located on chromosome IV, represented on the horizontal axis of Figure 6A. The distribution of the mean gene copy number per isolate, grouped by genetic cluster and separated by subgenome in allotriploids is summarized in Fig. 6B.

**Fig. 6.**
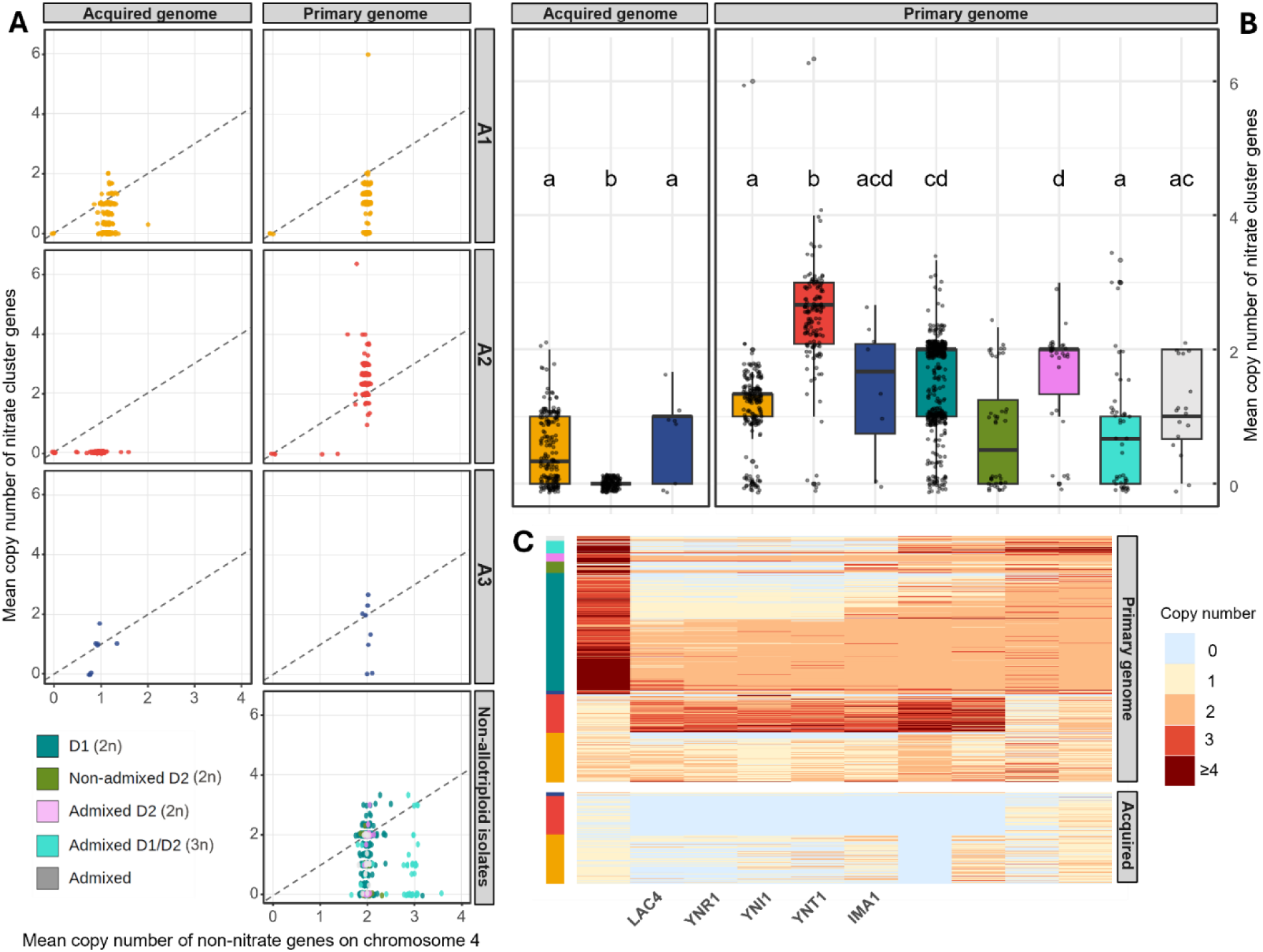
Copy-number variation (CNV) of nitrate assimilation genes relative to chromosome IV in *B. bruxellensis*. **(A)** Scatter plots showing, for each isolate, the relationship between the mean copy number of non-nitrate genes located on chromosome IV (x-axis) and the mean copy number of nitrate assimilation genes (y-axis). Each point represents one isolate, coloured according to its genetic cluster. For allotriploid clusters (A1, A2 and A3), primary genome and acquired genome are shown separately. The dashed diagonal line represents the expectation under uniform copy number between nitrate and non-nitrate genes. **(B)** Distribution of the mean copy number of nitrate assimilation genes across genetic clusters, shown as boxplots. Each box represents the distribution of isolate-level mean copy numbers for a given genetic cluster and genome. For allotriploid clusters, primary and acquired genomes are shown separately. Individual isolate values are overlaid as jittered points. Letters above boxes indicate statistically homogeneous groups based on post-hoc Dunn tests following Kruskal–Wallis analyses (Benjamini–Hochberg correction). **(C)** Heatmap showing copy-number variation across the ten first genes located on chromosome IV, including the three nitrate assimilation genes (*YNR1*, *YNI1* and *YNT1*) and flanking genes *LAC4* and *IMA1*. Each row represents one isolate, grouped by genetic cluster, and each column corresponds to a gene ordered along chromosome IV. For allotriploid clusters, copy numbers are shown separately for the primary and acquired genomes. Colour intensity reflects the copy number per gene, with darker colours indicating higher copy numbers.

At the chromosome-wide scale, mean copy number values are consistent with the expected ploidy of each genetic cluster (diploid, autotriploid, and allotriploid 2n+1n configurations; Fig. 6A), indicating stable copy number at the chromosome level.

In contrast, nitrate assimilation genes exhibit distinct patterns of copy-number variation among clusters. When considering total copy number across the nitrate assimilation genes without distinguishing between primary and acquired genome assignments, clusters display heterogeneous distributions that cannot be explained solely by ploidy (Supplementary Fig. 1). Among non-allotriploid clusters, nitrate gene copy number ranges from 0 to 3 copies (Fig. 6A). Clusters D1 and Admixed D2 display median values of two copies per gene, although 191/451 (43%) of D1 isolates carry only a single copy. Non-admixed D2 and the autotriploid Admixed D1/D2 cluster exhibit lower medians (0.5 and 0.67 copies per gene, respectively).

In allotriploid clusters, total copy numbers also show substantial variation, with median mean copy numbers per gene of 1.67, 2.67, and 2.0 in clusters A1, A2, and A3, respectively. These medians reflect the total number of copies detected per isolate. Given the composite genomic structure of allotriploid isolates, we next examined copy numbers separately for the two subgenomes (the primary and acquired genomes are expected to display 2 and 1 copies of each gene, respectively). In cluster A1, the mean copy number per nitrate gene ranges from 0 to 2 in both the primary and acquired genomes, with medians of 1.33 and 0.33, respectively (Fig. 6B). In cluster A2, copy numbers range from 2 to 4 in the primary genome, with a median of 2.67, whereas both the mean and the median are equal to zero in the acquired genome. This observation reflects the absence of nitrate assimilation genes in the available A2 acquired genome references (10), preventing the pipeline from assigning reads to an acquired genome copy during the competitive mapping. In cluster A3, values range from 0 to 3 in the primary genome and from 0 to 2 in the acquired genome, with medians of 1.7 and 1.0, respectively.

To determine whether these patterns reflect coordinated CNV across the entire nitrate gene cluster, or instead result from gene-specific variations within the cluster, we examined copy number of the first ten genes on chromosome IV (Fig. 6C). Each row corresponds to a single isolate, grouped by genetic cluster, with primary and acquired genomes displayed separately for allotriploid isolates. Overall, in 748 out of 946 isolates (≈79%), the three nitrate assimilation genes *YNR1*, *YNI1* and *YNT1* share identical copy numbers within a given isolate. This coordinated pattern is especially prevalent in non-allotriploid clusters, where 77%, 84%, 82%, and 95% of isolates exhibit uniform copy numbers across the three nitrate genes in the autotriploid Admixed D1/D2 cluster and in the diploid Admixed D2, D1, and Non-admixed D2 clusters, respectively. In contrast, allotriploid clusters display more heterogeneous copy number patterns. Across isolates from cluster A2, the three nitrate assimilation genes were not detected in the acquired genome, together with the two flanking genes and two downstream genes, forming a contiguous block of seven genes without assigned copies. These genes show increased copy numbers, with all detected copies being assigned to the primary genome possibly due to the lack of assignable acquired reference sequence for this locus, as noted above. Among the 143 isolates in this genetic cluster, 100 (≈70%) harbor three copies, with some isolates exhibiting four or more copies, while the remaining 43 isolates carry two copies. Only 35% of A2 isolates exhibit identical copy numbers across the three nitrate assimilation genes. In allotriploid cluster A1, 98 out of 191 isolates (≈51%) lack *YNR1* and *YNT1* in the acquired genome, whereas these genes occur at one to three copies in the primary genome. *YNI1* is retained as a single copy in 145 out of 191 isolates (≈76%) in the primary genome and in 91 out of 191 isolates (≈48%) in the acquired genome. Overall, coordinated copy numbers across the three nitrate genes are observed in 54% of acquired genomes and 48% of primary genomes within cluster A1. Within the allotriploid cluster A3, all isolates shared an identical copy number for each of the three nitrate assimilation genes. In this cluster, 38% of isolates exhibited two copies of each gene on the primary genome, whereas 50% carried a single copy of each gene on the additional genome. Among the 748 isolates showing uniform copy numbers across the three nitrate assimilation genes, 494 (≈66%) also display the same copy number for the flanking genes *LAC4* and *IMA1*.

Overall, these results indicate that CNV at the nitrate assimilation locus is predominantly coordinated at the cluster level, often extending to the flanking genes *LAC4* and *IMA1*, suggesting that this region behaves as a coherent genomic module. However, the increased heterogeneity observed in allotriploid clusters suggests more complex patterns of variation, consistent with independent gene-specific copy number loss.

### 6. Functional integrity of nitrate assimilation genes

We assessed the predicted functional integrity of nitrate assimilation genes across genetic clusters. Variant effect annotations were used to estimate the number of functional gene copies per isolate, separately for the primary genome and, when present, for the acquired genome. For each gene, functionality was quantified as the proportion of predicted functional copies among copies present, based on loss-of-function (LOF) annotations (including stop-gained, frameshift and start-loss) aggregated at the gene level (Fig.7A, B). A gene copy was considered putatively non-functional when at least one LOF variant was detected within its sequence. This approach enabled comparisons of functional integrity patterns among genetic clusters and between primary and acquired genomes, both for nitrate assimilation genes and for flanking genes located on chromosome IV. In Fig. 7A, the proportion of predicted functional copies among copies present is shown for each of the nitrate and flanking genes, aggregated by genetic cluster. In the primary genome, functionality of the three nitrate assimilation genes (*YNR1, YNI1 YNT1*), is largely conserved across all genetic clusters, with values exceeding 96%. The Admixed cluster displays a lower predicted functional proportion for the *YNI1*; however, this observation should be interpreted with caution given the genetic heterogeneity of this cluster, the difficulty in defining it unambiguously, and its limited representation (17/946). In contrast, more heterogeneous patterns were observed in the acquired genomes of allotriploid isolates for which nitrate assimilation genes could be assigned to the acquired genome, namely clusters A1 and A3. In the acquired genome of cluster A1, *YNR1* and *YNI1* exhibit predicted functional proportions below 60% relative to the number of copies present, whereas no predicted loss of functionality was detected for *YNT1*. In the acquired genome of cluster A3, predicted functionality for *YNR1*, *YNI1*, and *YNT1* is globally very low, falling below 20% and reaching 0% for the nitrate transporter gene *YNT1*. The two flanking genes, *LAC4* and *IMA1*, also display reduced predicted functional proportions in this acquired genome, with values of 25% and 33%, respectively.

**Fig. 7.**
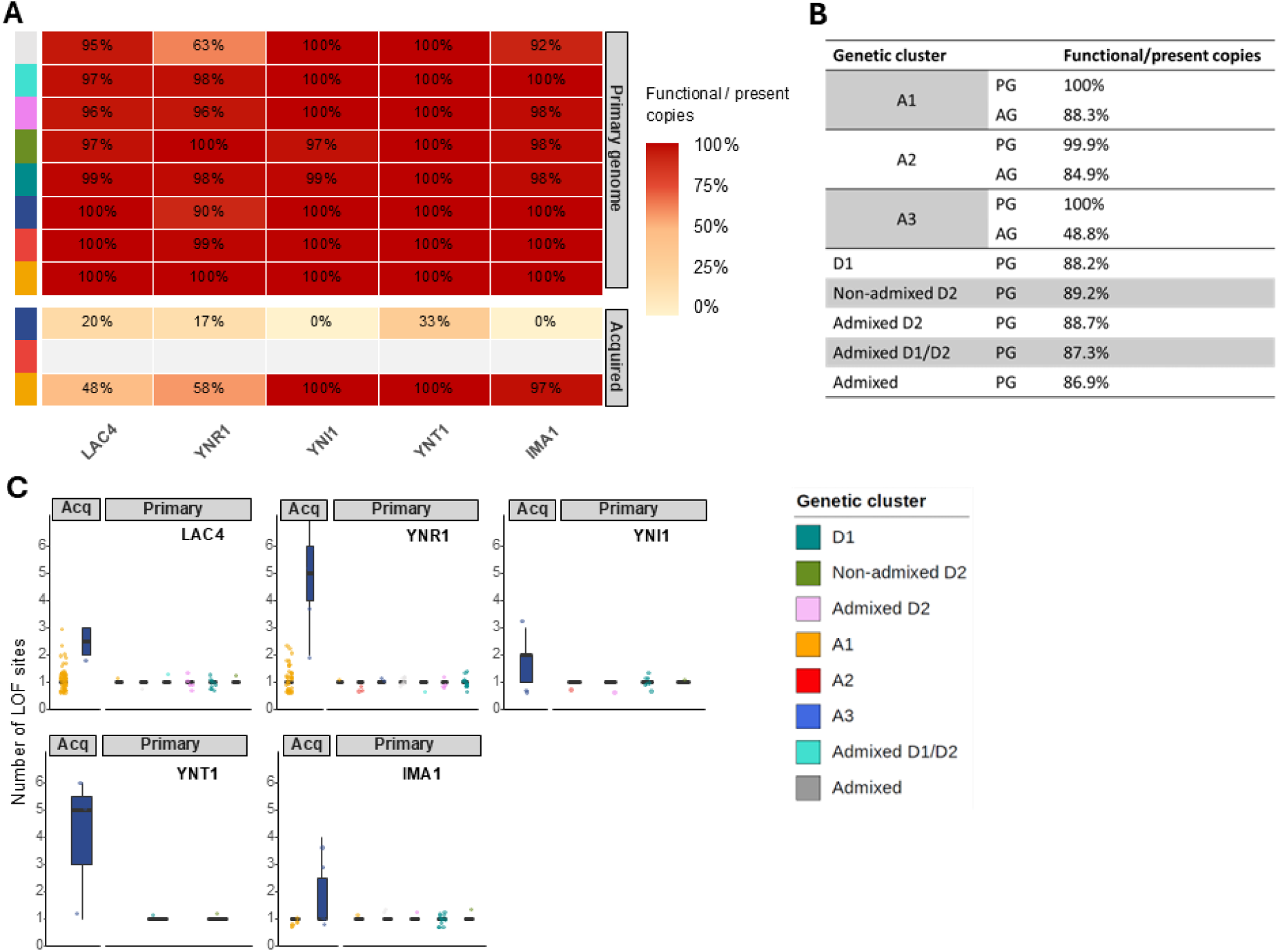
Functional integrity of genes on chromosome IV. **(A)** For each genetic cluster, the functional integrity of genes involved in nitrate assimilation (*YNR1, YNI1, YNT1*) and of two flanking genes (*LAC4* and *IMA1*) was quantified as the proportion of predicted functional copies among copies present (sum(fc) / sum(cp)). LOF variants include stop-gained, frameshift, start-loss and essential splice-site mutations. Values are shown separately for the primary genome and, when present, for the acquired genome. Percentages are indicated within each tile and represented using a color gradient from low (light) to high (dark) functional integrity. **(B)** Proportion of predicted functional gene copies among all gene copies present on chromosome IV for each genetic cluster. **(C)** Number of distinct LOF sites per isolate for each gene (*LAC4, YNR1, YNI1, YNT1* and *IMA1*), computed from variant effect annotations. Counts were obtained by aggregating distinct LOF positions per gene and per isolate. Values are shown separately for the primary and acquired genomes.

At the scale of the entire chromosome IV (Fig. 7B), the proportion of predicted functional genes in the primary genome remains consistently high across genetic clusters and never drops below 87%. In the acquired genomes of clusters A1 and A2, functional proportions of 88.3% and 84.9% were observed, respectively, values that are relatively high compared to those measured for nitrate assimilation genes in the acquired genome of cluster A1. In contrast, cluster A3 shows a markedly reduced predicted functional proportion at the chromosome IV scale, with only 48.8% of genes predicted to be functional. Although this value remains higher than those observed for nitrate assimilation genes in the acquired genome, it nevertheless indicates a comparatively low overall functional integrity in this acquired genome.

To further characterize the mutational basis underlying these differences in predicted functionality, we quantified the number of distinct LOF sites per gene and per isolate (Fig. 7C), considering only isolates carrying at least one LOF variant in the corresponding gene. Across all genetic backgrounds, stop-gained and frameshift variants represented the majority of predicted LOF mutations, both in primary and acquired genomes (Table S2). In the primary genome, LOF site counts remain low and constrained across all genes and genetic clusters, with most isolates carrying a single LOF site when present. This limited variation is consistent across *YNR1, YNI1, YNT1* and the flanking genes, with only occasional isolates exhibiting two or more sites. In contrast, acquired genomes display higher variability in LOF site counts. In cluster A1, isolates typically harbor one to two LOF sites per gene, especially in *LAC4* and *YNR1* genes, consistent with the reduced predicted functional proportions observed for these genes (Fig. 7A). In cluster A3, a higher mutational load is observed in the acquired genomes, particularly for *YNR1* and *YNT1*, where isolates frequently carry multiple distinct LOF sites per gene, with median values reaching up to five sites and broader distributions extending to higher values. Similar, although less pronounced, patterns are observed for *YNI1* and the flanking genes. Across all genes, this increased number of LOF sites in the acquired genome of A3 contrasts with the consistently low counts observed in the primary genome, indicating a clear difference in the accumulation of deleterious mutations between subgenomes. Notably, while LOF sites are generally limited to one per gene in the primary genome, the acquired genome of A3 frequently exhibits multiple independent LOF events within the same gene in a single isolate.

Consistent with the distribution of LOF site counts observed in Fig. 7C, LOF mutations displayed distinct positional patterns across genetic backgrounds (Fig. S2). In diploid and autotriploid backgrounds, LOF sites were generally isolate-specific and rarely shared among isolates (Fig. S2A). In contrast, acquired genomes of allotriploid clusters exhibited recurrent LOF positions shared across isolates within the same genetic cluster (Fig. S2B, Tab. S2). Cluster A3 additionally showed the co-occurrence of multiple recurrent LOF sites within the same genes and isolates.

Overall, these analyses reveal contrasted patterns of LOF accumulation across genetic backgrounds, ranging from rare and largely isolate-specific LOF events in diploid and autotriploid genomes to recurrent and multi-site LOF configurations in the acquired genomes of allotriploid clusters.

### 7. Nitrate utilization depends on the dosage and functional integrity of nitrate assimilation genes

To investigate the relationship between genetic variation at the nitrate assimilation locus and nitrate utilization, we then focused on the subset of 151 isolates for which growth on minimal medium with nitrate as the sole nitrogen source was experimentally assessed.

We first assessed whether nitrate utilization was associated with the presence of the complete nitrate assimilation gene cluster. Isolates were classified as carrying a complete cluster if at least one copy of all three genes were present (3/3), a partial cluster if one or two genes were present (1/3 or 2/3), and an absent cluster if none of the three genes were detected (Fig. 8A,B). Among isolates unable to utilize nitrate (NIT−), approximately half carried a complete nitrate gene cluster (n = 14). Among the remaining NIT− isolates, 12 lacked the entire cluster, while 2 carried a partial cluster comprising a single gene. In nitrate-assimilating isolates (NIT+), 77 isolates (≈89%) carried a complete cluster and the remaining 10 isolates carried a partial cluster (including 4 isolates with two genes and 6 with a single gene).

**Fig. 8.**
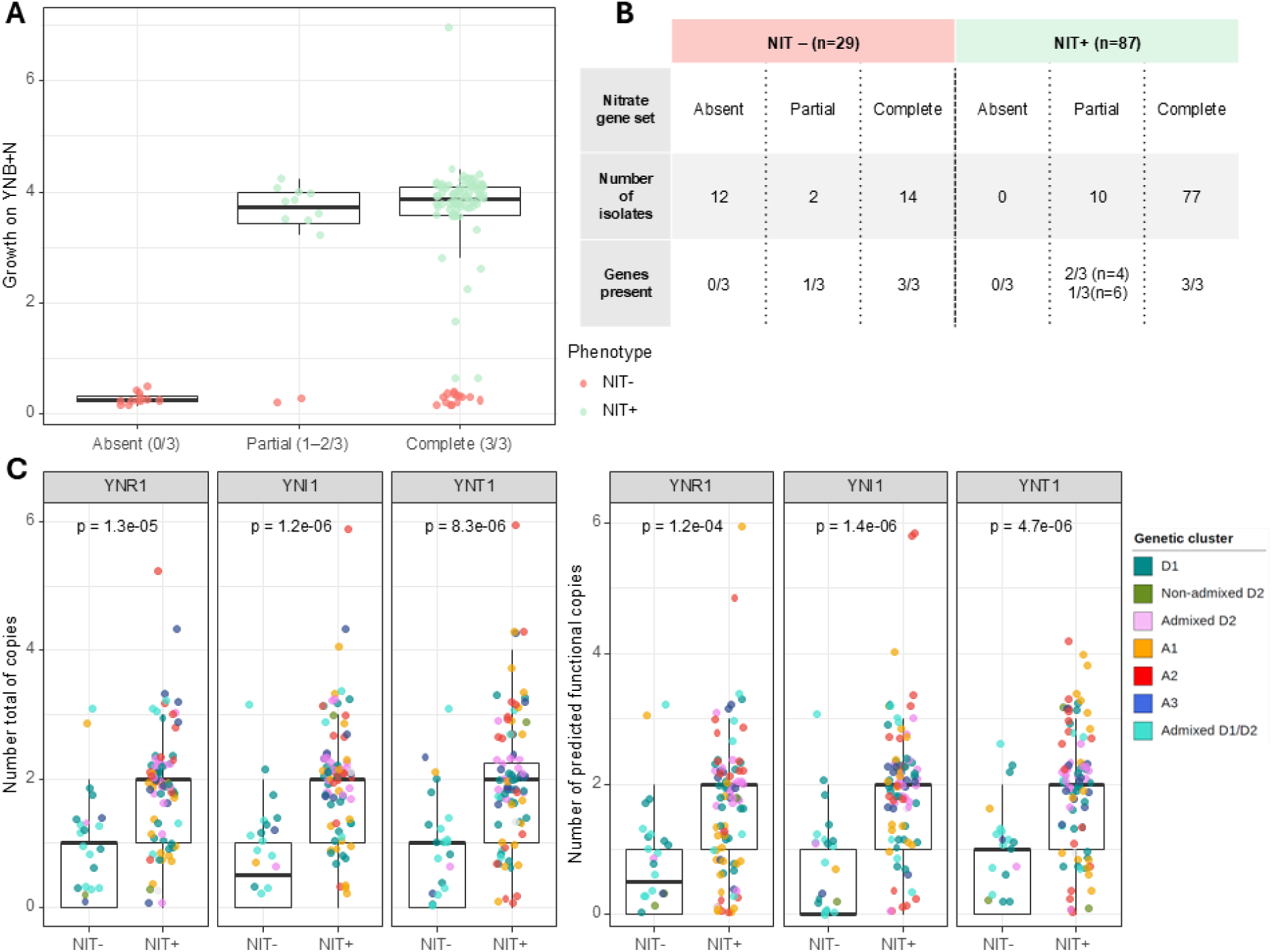
Genotype-phenotype relationships at the nitrate assimilation locus in 142 phenotyped subset. **(A)** Growth (maximal optical density) reached on nitrate (YNB+N minimum medium) of phenotyped isolates classified according to the completeness of the nitrate assimilation gene set (*YNR1, YNI1, YNT1*): absent (0/3 genes present), partial (1–2/3 genes present), and complete (3/3 genes present). Individual data points are shown, colored by phenotype (NIT− in pink, NIT+ in green. **(B)** Contingency table summarizing the distribution of NIT− and NIT+ isolates according to nitrate gene set completeness. **(C)** Distribution of copy number (left panels) and predicted functional copy number (right panels) for each nitrate assimilation gene (*YNR1, YNI1, YNT1*), stratified by phenotype (NIT− vs NIT+). Points represent individual isolates and are colored according to genetic cluster. Statistical differences between phenotypes were assessed independently for each gene using two-sided Wilcoxon rank-sum tests.

We next examined whether variation in copy number of the three nitrate assimilation genes, beyond simple presence or absence, was associated with nitrate utilization. Gene-specific copy number distributions were analyzed according to phenotype (NIT⁺ vs NIT⁻), distinguishing between total copy number (left panels) and copies predicted to be functional based on LOF annotations (right panels) (Fig. 8C). For all three genes, *YNR1*, *YNI1*, and *YNT1*, NIT+ isolates displayed significantly higher copy numbers than NIT− isolates, with median copy number reached two copies per gene, both when considering all copies present and when restricting the analysis to predicted functional copies. In contrast, among NIT− isolates, median copy numbers were lower and differed between total and predicted functional copies. For *YNR1*, the median number of copies present was 1, decreasing to 0.5 predicted functional copies. For *YNI1*, the median was 0.5 copies present and zero predicted functional copies. For *YNT1*, median values for both total and functional copies were equal to 1. No specific patterns associated with genetic cluster or ploidy were observed for these copy-number distributions.

The relationship between growth on nitrate, measured as the maximal optical density reached, and the mean copy number of the three nitrate assimilation genes was examined across genetic clusters (Fig. 9A). Among allotriploid isolates (left panel), including clusters A1, A2, and A3, 9/50 isolates are able to grow on nitrate with a mean copy number of one for the three nitrate assimilation genes. These isolates are predominantly from cluster A1 (6/9). In contrast, isolates from clusters A2 and A3 generally carry two or three copies of these genes when exhibiting growth on nitrate. Among non-allotriploid genetic clusters (right panel), isolates with a mean copy number of zero are predominantly unable to grow on nitrate. Isolates with a mean of one copy per gene show greater variability in growth, but the median growth remains low (≈0.39 OD), indicating that most isolates are unable to grow on nitrate. In contrast, isolates carrying two or three copies of the nitrate assimilation genes predominantly show growth on nitrate-containing minimal medium, a pattern notably observed in the Admixed D2 and D1 clusters. The few autotriploid Admixed D1/D2 isolates that successfully grew on nitrate exhibited a mean copy number of three for each of the three genes.

**Fig. 9.**
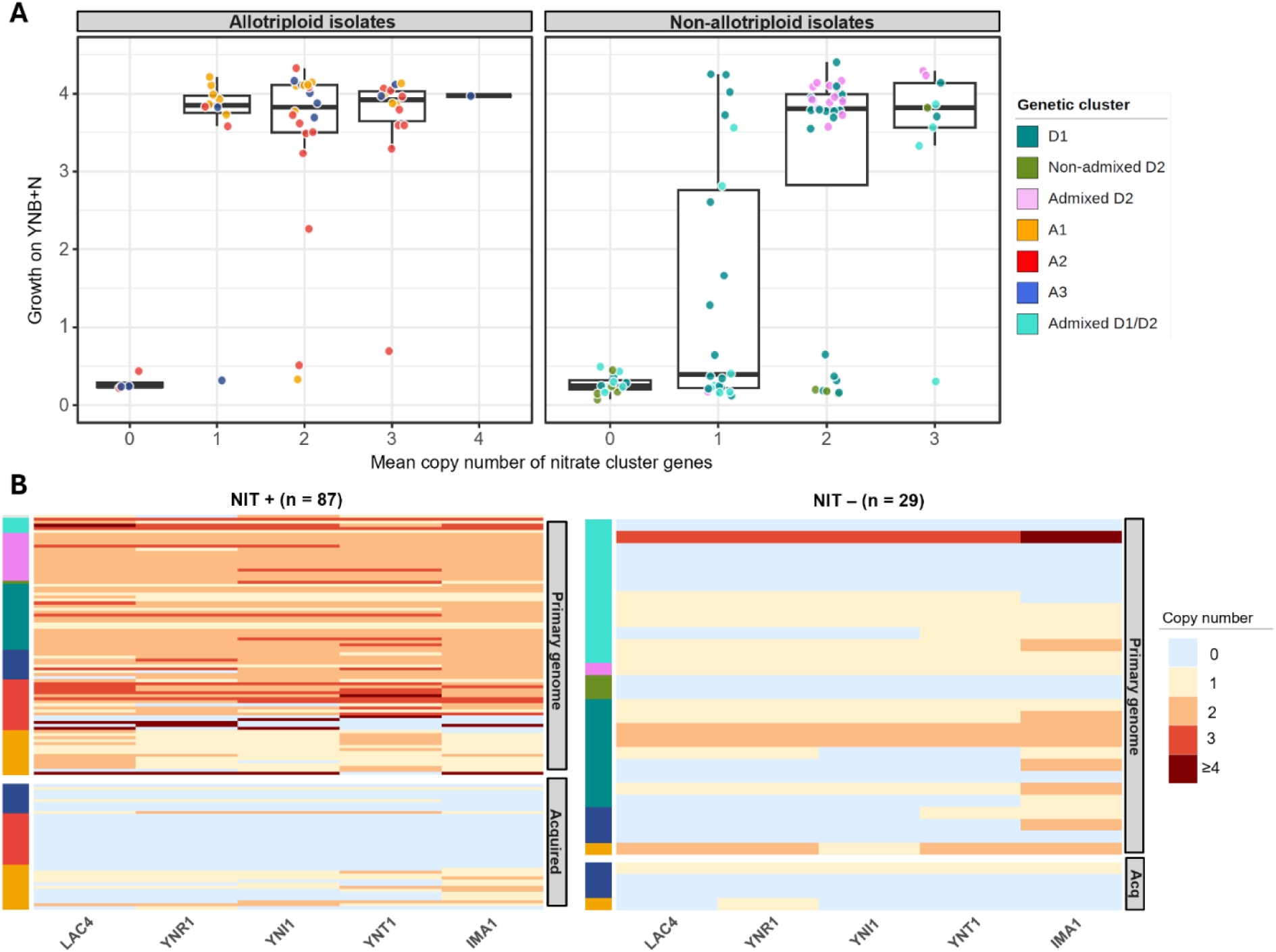
Relationship between nitrate gene copy number and nitrate assimilation phenotype. **(A)** Growth on nitrate (YNB+N minimum medium) plotted against the mean copy number of nitrate assimilation genes (YNR1, YNI1, YNT1) per isolate. The mean copy number was calculated as the average number of copies per gene across the three-gene nitrate cluster. Allotriploid isolates (A1, A2, A3) and non-allotriploid isolates are shown in separate panels. Individual isolates are represented as points colored according to their genetic cluster. **(B)** Heatmaps showing copy-number variation of nitrate assimilation genes (*YNR1, YNI1, YNT1*) and flanking genes (LAC4 and IMA1) on chromosome IV in NIT+ (left, n = 87) and NIT− (right, n = 29) isolates. Copy number is shown separately for the primary genome and, when present, the acquired genome in allotriploid isolates. Rows correspond to individual isolates, ordered by genetic cluster, and colors indicate copy number from low (light) to high (dark).

The distribution of gene copies between the primary and acquired genomes was examined among nitrate-assimilating (NIT⁺) allotriploid isolates (Fig. 9B). Within the cohort of 87 NIT+ isolates retained for the analysis, cluster A2 displays patterns representative of those observed in the full dataset of 946 isolates, with nitrate assimilation genes detected in three or more copies and assigned only to the primary genome, possibly reflecting the lack of assignable acquired reference sequence. In this cluster, some isolates lack one or two of the nitrate assimilation genes, while other genes of the same cluster are present at four or more copies. In cluster A1, the primary genome most often carries a single copy of each nitrate assimilation gene, while copies are frequently absent from the acquired genome. When the genes are not detected in the acquired genome, they are typically present as one or two copies in the primary genome. Conversely, when a single copy is present in the acquired genome, a corresponding single copy is generally also observed in the primary genome. In a small number of cases, two copies of these genes were detected in the acquired genome within this cluster. In cluster A3, the primary genome predominantly carries two copies of the nitrate assimilation genes, whereas the acquired genome frequently carries one copy and more often none. Finally, among the 29 nitrate non-assimilating (NIT−) isolates, the majority carry either zero or a single copy of each of the three nitrate assimilation genes.

## Discussion

### Functional gene dosage as the main determinant of nitrate assimilation

Our results indicate that nitrate assimilation in *B. bruxellensis* is primarily determined by functional gene dosage rather than by the sole presence or absence of the genes involved. The discordance between gene presence and nitrate assimilation, together with the higher functional copy number observed in NIT+ isolates, supports nitrate assimilation as a dosage-dependent quantitative trait. Across most genetic backgrounds, nitrate assimilation is associated with approximately two functional copies per gene, suggesting a minimal functional dosage required for effective assimilation (32,33).

This dosage requirement is consistent with the metabolic structure of the pathway. Nitrate assimilation involves active transport followed by sequential reduction of nitrate to nitrite and ammonium, reactions that require coordinated enzymatic activity and substantial reducing power (27,28). Insufficient gene dosage may therefore limit the pathway activity even when the genes themselves are present. However, the relationship between copy number and phenotype is not strictly fixed across genetic backgrounds. In particular, some isolates from the allotriploid A1 cluster grow on nitrate despite carrying a single apparent copy per gene, suggesting that regulatory context or broader genomic background may modulate the effective dosage required for pathway activity. Conversely, autotriploid isolates tend to exhibit nitrate assimilation at higher gene dosages, typically close to three copies per gene, consistent with their ploidy level. Given that most tested isolates in this cluster harbor fewer than one copy per gene on average, this higher dosage requirement likely contributes to the rarity of the NIT+ phenotype in the autotriploid cluster.

Beyond copy number, predicted loss-of-function mutations further reduce the functional dosage of the pathway, particularly within acquired genomes of allotriploid isolates. Although functional status inferred from predicted loss-of-function annotations remains an approximation, this potential bias is applied uniformly across all strains. Overall, gene dosage and predicted functional integrity therefore act jointly to shape nitrate assimilation, supporting this phenotype as a quantitative metabolic trait governed by functional gene dosage.

### Nitrate assimilation as an ancestral trait under relaxed selection in specific lineages

At the species scale, our results confirm and refine previous reports of pronounced heterogeneity in nitrate assimilation within *B. bruxellensis* (16,20,21,25,26). Nitrate assimilation is strongly structured by the genetic clustering of the species, yet nitrate-assimilating isolates are detected in every genetic cluster of the phenotyped dataset. This broad yet uneven distribution suggests that nitrate assimilation may represent an ancestral trait of the species that has subsequently followed heterogeneous evolutionary trajectories among lineages. Copy-number reduction, locus loss, and predicted loss-of-function mutations reduce the effective functional dosage of the pathway in several clusters. Moreover, the contrasting distribution of LOF mutations, ranging from predominantly isolate-specific events in diploid and autotriploid backgrounds to recurrent and shared mutations in allotriploid acquired genomes, suggests lineage-and subgenome-specific histories of functional erosion. Such processes are commonly associated with relaxation of selective constraints on metabolic pathways (34,35), and our results therefore likely reflect differences in selective constraints across lineages.

The ecological environments in which *B. bruxellensis* occurs may contribute to these heterogeneous evolutionary trajectories. This yeast is primarily associated with anthropized fermentation ecosystems such as wine, beer, kombucha, and bioethanol production, characterized by fluctuations in nutrient availability, microbial competition and harsh physicochemical conditions including elevated ethanol concentrations (7,15,36). Although genetic clusters are more frequently associated with one specific ecological niche (Fig. 10), this correspondence remains imperfect, as strains may circulate between fermentation environments and their isolation source may not reflect their full ecological history (21). Nevertheless, differences in nitrogen composition among fermentation environments may contribute to the uneven distribution of nitrate assimilation across lineages. Although quantitative data remain limited, available studies suggest that nitrate concentrations tend to be higher in brewing environments than in wine fermentations (37,38). Consistently, the A1 cluster, frequently associated with brewery environments, tends to display a higher prevalence of nitrate-assimilating isolates, whereas clusters more commonly isolated from wine environments, notably the autotriploid Admixed D1/D2 and diploid D1 clusters, tend to contain a comparatively larger proportion of NIT-strains. Interestingly, *IMA1*, one of the genes flanking the nitrate assimilation cluster, encodes an isomaltase involved in α-glucoside utilization in *S. cerevisiae* (39). Given the relevance of these sugars in brewing environments and the association of subtelomeric sugar-utilization gene with yeast ecological lifestyles, the proximity of *IMA1* to nitrate assimilation genes raises the possibility that this subtelomeric region experiences selective constraints related to both carbon and nitrogen utilization (39). However, no clear A1-specific pattern of functional variation in *IMA1* was observed in our dataset. Overall, these ecological patterns are consistent with potential differences in selective constraints acting on the nitrate assimilation pathway. In nitrate-rich environments, loss of functional nitrate assimilation genes may be more likely to be counter-selected, contributing to the maintenance of functional gene copies. Conversely, where nitrate is scarce and other nitrogen sources are abundant, weaker selective constraints may allow gene loss or functional erosion to accumulate over evolutionary time. More broadly, ecological heterogeneity may also contribute to the variability of nitrogen utilization within the species. In addition to nitrate assimilation, a substantial proportion of isolates exhibit ammonium-related auxotrophies (AMMO−), despite ammonium being a preferred nitrogen source in many yeasts (40). The coexistence of NIT+, NIT−, and AMMO− phenotypes therefore indicates that nitrogen-use pathways have undergone substantial modification within the species.

**Fig. 10.**
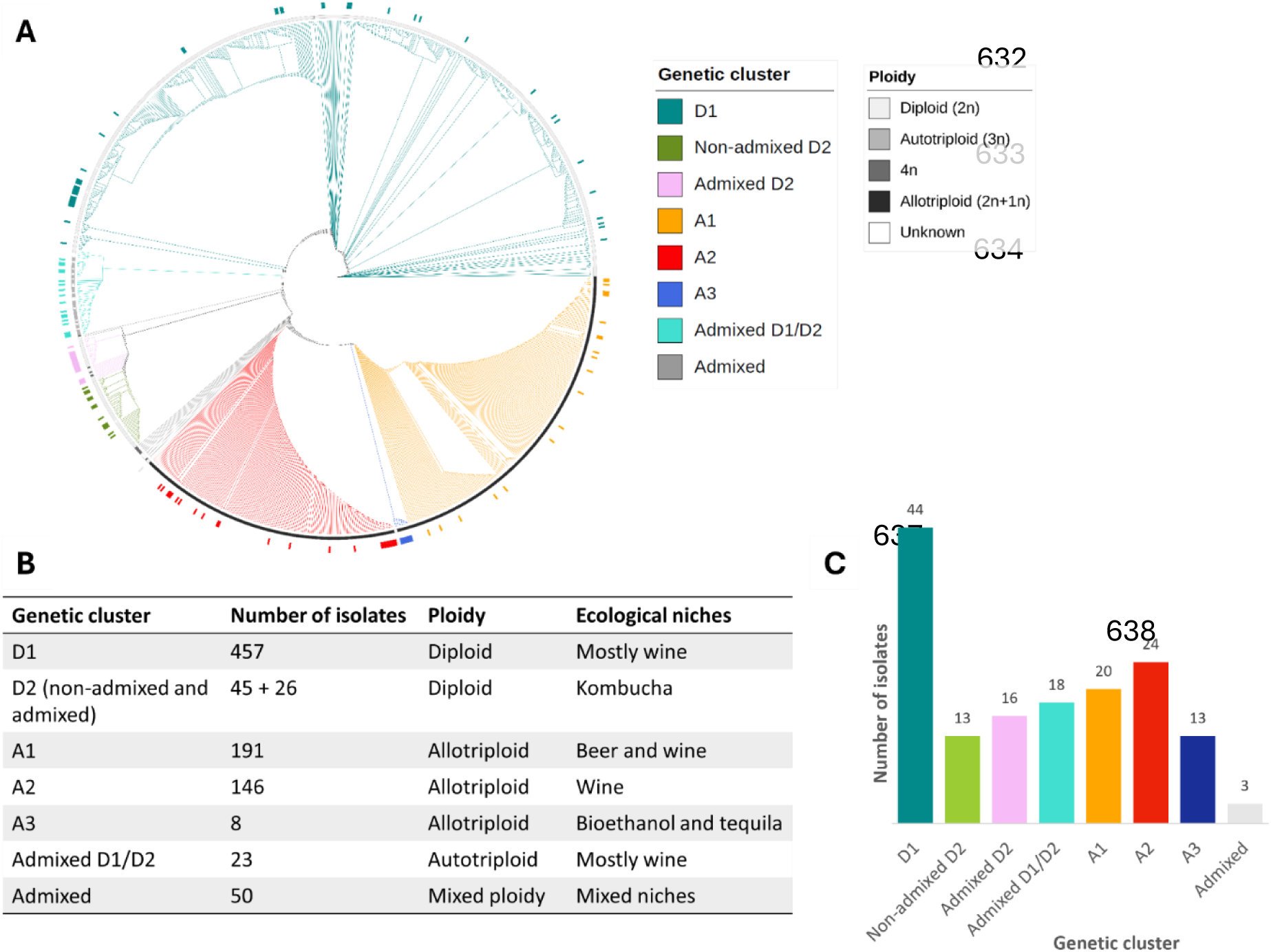
Genetic diversity, ploidy distribution and ecological origin of the *Brettanomyces bruxellensis* strains analyzed in this study. **(A)** Circular phylogenetic tree built from whole-genome sequences of the 946 sequenced isolates and visualized using iTOL (Letunic & Bork, 2024). Colored branches indicate the previously defined genetic clusters and admixed groups. The 151 strains selected for phenotypic analyses are highlighted on the tree using a color strip. Ploidy levels are represented as an outer grey-scale annotation ring. **(B)** Summary of the main characteristics of each genetic cluster, including number of isolates in the whole dataset, predominant ploidy level and major ecological niches of isolation. **(C)** Number of isolates included in the phenotypic dataset for each genetic cluster.

Together, these observations support the view that nitrate assimilation represents an ancestral trait that has undergone lineage-specific relaxation of selection, resulting in varying degrees of pathway preservation or loss.

### Polyploidy and genome architecture shape nitrate assimilation

In addition to ecological context, polyploidy is associated with a broader diversity of genomic configurations through which the functional dosage of nitrate assimilation genes may be maintained or reduced in *B. bruxellensis*. Increased ploidy introduces additional variation in gene copy number, allelic composition, and subgenome-specific contribution, resulting in multiple genomic architectures within the species (11,41). This effect is particularly pronounced in allotriploid lineages, where the coexistence of a primary genome and an acquired genome is associated with differential retention, amplification and functional degradation of nitrate assimilation genes across subgenomes.

The three allotriploid clusters illustrate different aspects of this genomic diversity. In A2, nitrate assimilation genes cannot be assigned to the acquired genome during competitive mapping, likely reflecting the absence of the corresponding loci in the available acquired genome references (10). Consequently, it remains unclear whether nitrate assimilation genes are truly absent from the acquired genome or simply not represented in the available references. Alternatively, this pattern could reflect either an ancestral absence of the nitrate assimilation cluster in the donor lineage or a secondary loss following hybridization (42). In these isolates, multiple intact copies of the nitrate assimilation genes are consequently detected and assigned to the primary genome, resulting in elevated functional dosage. Consistently, all phenotyped A2 isolates exhibit the NIT⁺ phenotype.

In A1, functional copies are distributed across both subgenomes or concentrated within the primary genome, but acquired-genome copies frequently carry predicted LOF mutations. A related but more pronounced configuration is observed in A3, where extensive functional degradation affects the acquired genome at the nitrate assimilation locus and more broadly across chromosome IV, leaving nitrate assimilation associated with copies retained in the primary genome. Autotriploid isolates represent a different genomic configuration, generally harboring fewer functional nitrate assimilation gene copies and displaying the lowest prevalence of the NIT+ phenotype.

Taken together, these observations indicate that polyploidy does not correspond to a single genomic configuration for nitrate assimilation. Instead, variation in ploidy and subgenome composition is associated with multiple genomic architectures through which functional gene dosage may be maintained, redistributed, or reduced. Such mechanisms may be particularly relevant in largely clonal microorganisms, where structural variation, gene dosage modulation, and genome rearrangements can contribute substantially to evolutionary change in the absence of frequent sexual recombination (43,44). In *B. bruxellensis*, hybridization and polyploidy may therefore contribute to the diversity of genomic architectures observed across lineages. Across allotriploid lineages, the primary genome appears to represent the main carrier of nitrate assimilation, whereas acquired genomes frequently exhibit greater functional degradation, including LOF accumulation and shared mutational hotspots. This asymmetry is consistent with previous observations showing a stronger contribution of the primary genome than the acquired genome to phenotypic variation in *B. bruxellensis* (11).

Altogether, our results show that nitrate assimilation in *B. bruxellensis* is best understood as a quantitative metabolic trait primarily governed by functional gene dosage and shaped by genome architecture. Variation in gene copy number, functional integrity, and subgenome composition generates multiple genomic configurations associated with nitrate assimilation across the species. These patterns are consistent with nitrate assimilation representing an ancestral metabolic ability whose evolutionary maintenance varies among lineages under differing ecological selective regimes.

## Materials and Methods

### Phenotypic analysis

#### 1. B. bruxellensis strains

We selected 151 *B. bruxellensis* strains representative of the genetic diversity of the species, belonging to the seven previously described genetic lineages, as well as admixed strains that do not cluster within any distinct lineage (10,11). These strains are highlighted on the phylogenetic tree of the 946 isolates for which whole-genome sequences are available (Fig. 10A; (11)). Strain names are provided in the supplementary data. The ecological niches shown in Fig. 10B correspond to the predominant niches associated with each genetic cluster, although some isolates may have been recovered from other environments.

#### 2. Media and growth conditions

Yeasts were first grown in YPD liquid medium (yeast extract 10 g·L⁻¹, peptone 20 g·L⁻¹, dextrose 20 g·L⁻¹) for 48 h at 24 °C without shaking. Cells were then washed in sterile water, and viable cell concentration was assessed using a flow cytometry approach (CytoFLEX, Beckman Coulter, Brea, CA, USA). Fifty microliters of the homogenized culture were mixed with 3 µL of propidium iodide (PI), diluted to a final volume of 1 mL in McIlvaine buffer (0.2 M sodium phosphate dibasic, 0.1 M citric acid), and analyzed by flow cytometry (45)(Zimmer et al., 2014). Fifty thousand events were recorded per experiment. Cell viability was estimated using 488 nm excitation source with a 610/20 nm bandpass filter to detect PI fluorescence (dead cells). Growth assays were performed in liquid synthetic medium containing 1.7 g·L⁻¹ YNB w/o (yeast nitrogen base without ammonium and amino acids) (DIFCO, Thermo Fisher Scientific, MA, USA), 20 g·L⁻¹ D-glucose, and supplemented either with (i) 0.66 g·L⁻¹ ammonium sulfate to obtain 5 mM NH₄⁺ for the ammonium condition (YNB+A), or (ii) 0.4 g·L⁻¹ sodium nitrate to obtain 5 mM NO₃⁻ for the nitrate condition (YNB+N). Growth in YPD liquid medium was used as a control. Growth was monitored in 96-well microplates at 24 °C, with 200 µL of medium per well under agitation (CLEARLine, Thermo Fisher Scientific, MA, USA), and optical density was measured every two hours for 96 h at 600 nm using a spectrophotometer (FLUOstar Omega, BMG LABTECH). Each strain was inoculated at 2.5 × 10⁵ viable cells·mL⁻¹ in triplicate for each medium.

#### 3. Determination of growth parameters

Growth curves were obtained for each strain and each medium in triplicates. Growth curves were analyzed using generalized additive models (GAMs), a nonparametric regression framework allowing flexible modeling of growth dynamics without assuming a predefined parametric form (46,47). GAMs were fitted with optical density as a function of time, using a smooth spline function, allowing flexible modeling of growth trajectories. From the GAM-fitted curves, optical density values were predicted continuously over time. Growth parameters were then extracted from these predictions (48). The instantaneous growth rate, r(t), was computed as the numerical derivative of the fitted curve with respect to time. The maximum instantaneous growth rate (rₘₐₓ) was retained as a proxy for growth velocity. The maximum population size (K) was defined as the maximum predicted optical density (OD_max) reached during the growth. Lag phase duration was estimated only for strains exhibiting measurable growth (OD_max > 0.5) and was defined as the time required to reach 10% of OD_max. All growth parameters were therefore derived from GAM-based predictions, ensuring consistency across strains and conditions and minimizing the influence of experimental noise on parameter estimation. For the determination of nitrogen source utilization, a strain was considered able to grow when the final population reached an optical density greater than 0.5.

#### 4. Statistical analysis

Statistical analysis were performed to investigate both the distribution of qualitative phenotypes (NIT+, NIT-, AMMO-classes) and the quantitative variation of growth parameters across genetic populations and nitrogen conditions. Associations between phenotypic classes and genetic populations were first assessed using contingency table analyses and tested using Pearson’s chi-squared (χ²) test. To identify which population–phenotype combinations contributed most strongly to significant associations, standardized residuals were computed and adjusted following Haberman’s correction, yielding approximately normally distributed Z-scores for each cell. Two-tailed p-values derived from these residuals were corrected for multiple testing using the Benjamini–Hochberg procedure to control the false discovery rate (FDR).

Quantitative growth parameters (maximum population size K, maximum growth rate rₘₐₓ and lag phase duration) extracted from smoothed GAM-fitted curves were analyzed using linear mixed-effects models (LMMs). For each parameter, the following model was fitted :

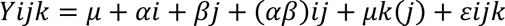

where Yijk represents the growth parameter measured for strain *k* belonging to genetic population *j* in nitrogen condition *i*; μ is the global intercept; α_i_ is the fixed effect of nitrogen condition; β_j_ is the fixed effect of genetic population; (αβ)_ij_ is the interaction between nitrogen condition and population; u_k(j)_ is a random intercept for strain *k* nested within population *j*, accounting for repeated measurements and inter-strain variability; and ε_ijk_ is the residual error term.

Fixed effects were tested using type III analysis of variance (ANOVA) on the mixed-effects models, with degrees of freedom adjusted according to the Kenward–Roger approximation, providing robust F-tests in the presence of unbalanced designs and random effects. Post hoc pairwise comparisons were performed on estimated marginal means using Tukey’s honestly significant difference (HSD) procedure, with Kenward–Roger–adjusted degrees of freedom.

Model assumptions were evaluated by inspecting residual normality, homoscedasticity, and the distribution of random effects. When necessary, response variables were log-transformed to stabilize variances and improve model fit. All statistical analyses were conducted in R using agricolae and emmeans packages for mixed-effects modeling and multivariate analyses (49).

#### 5. *In silico* pairwise growth simulation

*In silico* pairwise growth simulations were performed to estimate the relative fitness of strains based on their growth characteristics measured in monoculture. This approach integrates experimentally derived monoculture growth parameters into a deterministic growth model to simulate pairwise population dynamics under no-interaction assumption (50,51). These simulations do not model direct biological interactions between strains. Instead, they provide a deterministic approximation of relative population expansion based solely on independently measured monoculture growth dynamics. Consequently, the simulated outcomes should be interpreted as model-based estimates of relative growth contribution rather than direct measurements of competitive fitness.

For each strain and growth condition, monoculture growth curves were fitted using generalized additive models (GAMs), as described above, providing smooth estimates of population over time. The predicted growth trajectory for strain *i* in a given medium is denoted Ni(t), corresponding to the GAM-predicted optical density at time *t*.

For each pair of strains A and B grown under the same condition, their predicted growth curves N_A_(t) and N_B_(t) were evaluated on a common time grid spanning the observed experimental time range. At each time point, the relative contribution of each strain to the total predicted population was calculated as:

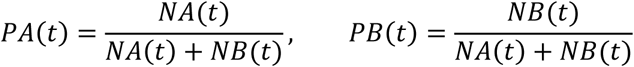

*In silico* simulation was considered as completed when the maximal population was reached. The predicted outcome of each simulation was defined as the final relative fraction of strain A, denoted PA(t_final_), and was expressed as a percentage. Values around 50% indicate equal colonization of the medium by both strains, and values approaching 0% of 100% indicate that one strain was fully outcompeted by the other. This procedure was applied to all pairwise strain combinations within each growth condition, yielding strain-by-strain matrices of *in silico* competitions.

For each strain and medium, mean predicted final relative population was calculated as the average final population percentage obtained across all pairwise simulations. Distributions of these values were compared among genetic populations using non-parametric Kruskal– Wallis tests.

### Genetic analysis

#### 1. Whole-genome sequences data and reference genomes

Whole-genome sequencing data were obtained for 1,060 isolates selected to be representative of the *B. bruxellensis* whole species. Ecological and geographical metadata, as well as sequencing protocols for these isolates, are described in Loegler et al., 2025 (11).

All sequencing data are publicly available in the European Nucleotide Archive (ENA) under accession numbers PRJEB28420, PRJEB71961, and PRJEB41126. Raw sequencing data were retrieved as paired-end FASTQ files (R1/R2) for each isolate. Reads were first subjected to a trimming step using fastp v1.0.1 to remove sequencing adapter remnants and filter out short reads (52). Following an additional quality control step, 1,023 isolates were retained for downstream analyses. The detailed pipeline described below was applied to chromosome IV of the *B. bruxellensis* genome, which harbors the genes of the nitrate assimilation cluster. All genomic analyses were conducted using the annotated diploid reference genome UMY321 as the primary reference (53). In addition, three haploid assemblies representing the acquired genomes, described in Eberlein et al., 2021 (10), were used: A1 (formerly Beer), A2 (formerly Wine 1), and A3 (formerly Teq/EtOH).

#### 2. Reads mapping

All sequencing reads from all isolates were first mapped to the diploid reference genome of *B. bruxellensis* UMY321 (53). Read alignment was performed using the BWA-MEM2 algorithm implemented in BWA v0.7.17-r1188 (54). Duplicate reads were marked, and reads with a mapping quality below 30 were discarded using samtools v1.21 (55). Ploidy information for each isolate was retrieved from Loegler et al., 2025 (11) and used in subsequent analyses.

#### 3. Primary and acquired genome detection and read separation

For this analysis, we applied a methodology similar to that previously described by Loegler et al., 2025 (11). Briefly, in addition to mapping all reads to the diploid reference genome, we performed competitive mapping of reads from allotriploid isolates against primary and acquired genomes, using the BWA-MEM2 algorithm implemented in BWA v0.7.17-r1188 (54). To this end, concatenated reference genomes were generated for each acquired genome (A1, A2, and A3) by combining the diploid *B. bruxellensis* reference genome with each of the assembled acquired genomes (Beer, Wine 1, and Teq/EtOH) from Eberlein et al., 2021 (10).

Reads from each allotriploid isolate were independently mapped against each of the three concatenated references. As described by Loegler et al.; 2025 (11), an acquired genome was considered present when the proportion of high-quality mapped reads assigned to the acquired genome exceeded 11.5%. This cutoff was defined based on previously available data from *B. bruxellensis* allotriploid isolates (11).When more than one acquired genome exceeded this threshold, the acquired genome with the highest proportion of high-quality mapped reads was retained as the major acquired genome. Ambiguously mapped reads (MAPQ<20) were excluded to ensure strict genome assignment during competitive mapping, acknowledging that the treatment of multimappers can influence downstream analyses (56). This filtering step resulted in the removal of only 13.2% of reads on average across allotriploid isolates. Following this filtering step, reads were assigned either to the primary genome or to one of the acquired genomes based on their mapping results. Reads assigned to the primary genome were subsequently remapped to the diploid reference genome, while reads assigned to each acquired genome were remapped to their corresponding assembled reference genomes and, in parallel, to the diploid reference genome to allow direct and consistent comparisons between primary and acquired genomes.

#### 4. CNV calling

To infer gene copy number for each isolate on the diploid primary genome and, when relevant, on acquired genomes, sequencing coverage was estimated using mosdepth v0.3.11 (57), which computes read depth over predefined genomic intervals corresponding to gene coordinates. Analyses explicitly accounted for isolate ploidy, such that 100% coverage corresponded to the expected coverage of a gene present at the copy number predicted by ploidy. Based on the results reported by Loegler et al., 2025 (11), ploidy was set to three for isolates belonging to the autotriploid (3n) cluster, to two for diploid isolates as well as for the primary genome (PG) of allotriploid isolates, while acquired genomes (AG) in allotriploids were considered haploid. To ensure robust copy number inference across mapping strategies, a breadth-of-coverage threshold of 10× was applied for diploid and autotriploid (3n) genomes mapped to the diploid reference genome, whereas a breadth threshold of 1× was used for allotriploid PG and AG mapped to concatenated reference genomes. Within the subset of 151 phenotyped strains, nine isolates were excluded because their sequencing data did not allow reliable gene copy-number estimation using our bioinformatic pipeline (YJS5397, YJS5454, YJS5459, YJS5469, YJS7831, YJS7858, YJS8072, YJS8229, YJS8285).

#### 5. SNP calling

Variant calling was performed using bcftools v1.22, applying the mpileup and call functions to files obtained from the different mapping strategies (58). First, SNP calling was conducted on BAM files resulting from the mapping of diploid and autotriploid isolates to the *B. bruxellensis* diploid reference genome (53), as well as on reads assigned to the primary genome during competitive mapping and subsequently remapped to this same reference. Second, SNP calling was performed on reads assigned to each acquired genome following competitive mapping and remapped to the corresponding acquired genome assembly (10), as well as to the diploid reference genome to ensure the comparability between acquired and primary genomes. Ploidy parameters for the call command were set as described for CNV inference, with acquired genome calls treated as haploid and all other calls treated as diploid. The diploid reference genome was used together with its associated gene annotation. Assemblies of the acquired genomes were annotated using the annotation transfer tool Liftoff v1.6.3.2 (59), with the *B. bruxellensis* reference genome annotation used as the reference annotation set. In addition, reads assigned to the acquired genomes and remapped to the diploid reference were also annotated using the reference genome annotation, in order to complement the annotation transfer and to ensure maximal comparability between the primary genome and the acquired genomes. Individual variant calls were then merged per group for downstream analyses. Quality filters identical to those described in Loegler et al., 2025 (11) were subsequently applied to variant calls using bcftools v1.22. For inclusion in downstream analyses, allopolyploid isolates were required to pass all quality criteria at the whole-genome, primary-genome, and acquired-genome levels, resulting in 354 allopolyploid isolates retained. Non-allopolyploid isolates were required to pass whole-genome filters only, yielding a final dataset of 946 isolates out of the 1,060 sequenced isolates.

#### 6. Determination of gene functionality

To assess gene loss-of-functionality (LOF), functional impact annotations were obtained using the localized mode of the *BCFtools/csq* plugin implemented in bcftools v1.22 (60), applied to all variant call datasets. Predicted loss-of-function variants were used as proxies for gene functionality, acknowledging that such variants do not necessarily result in complete functional inactivation in all cases. Loss-of-function variants were defined as variants predicted to introduce stop-gained mutations, frameshifts, start-loss events, or disruptions of essential splice sites, as identified by *BCFtools/csq*. Variant-level annotations were subsequently aggregated at the gene level and analyzed separately for each gene copy in the primary genome (PG) and in the acquired genomes (AG) of allotriploid isolates. A gene copy was considered as putatively non-functional when at least one predicted loss-of-function (LOF) variant was detected within the corresponding gene sequence. However, these annotations should be interpreted as proxies for potential functional alteration rather than definitive evidence of complete gene inactivation, as the actual impact of predicted LOF variants may depend on their genomic context, position within the coding sequence, compensatory mechanisms, or regulatory effects (61).

## Author contributions

Audrey Vigna: Data curation, Formal analysis, Investigation, Methodology, Software, Visualization, Writing – original draft, Writing – review & editing

Jules Harrouard: Manipulation, Formal analysis, Conceptualization, Visualization, Methodology, Writing – review & editing

Cecile Miot-Sertier: Manipulation, Writing – review & editing

Louis Michelizza: Manipulation

Victor Loegler: Formal analysis, Resources, Writing – review & editing

Philippe Marullo: Methodology, Writing – review & editing

Anne Friedrich: Conceptualization, Methodology, Writing – review & editing

Joseph Schacherer: Conceptualization, Methodology, Writing – review & editing

Emilien Peltier: Conceptualization, Formal analysis, Visualization, Methodology, Writing – review & editing

Warren Albertin; Conceptualization, Investigation, Formal analysis, Methodology, Writing – original draft, Writing – review & editing

## Acknowledgments

The authors thank the GPR Bordeaux Plant Sciences-DYNASTY consortium for insightful discussions (WP8-DYNASTY received financial support from the French government in the framework of the IdEX Bordeaux University “Investments for the Future” program/GPR Bordeaux Plant Sciences).

## Supplementary data

**Supp.1.**
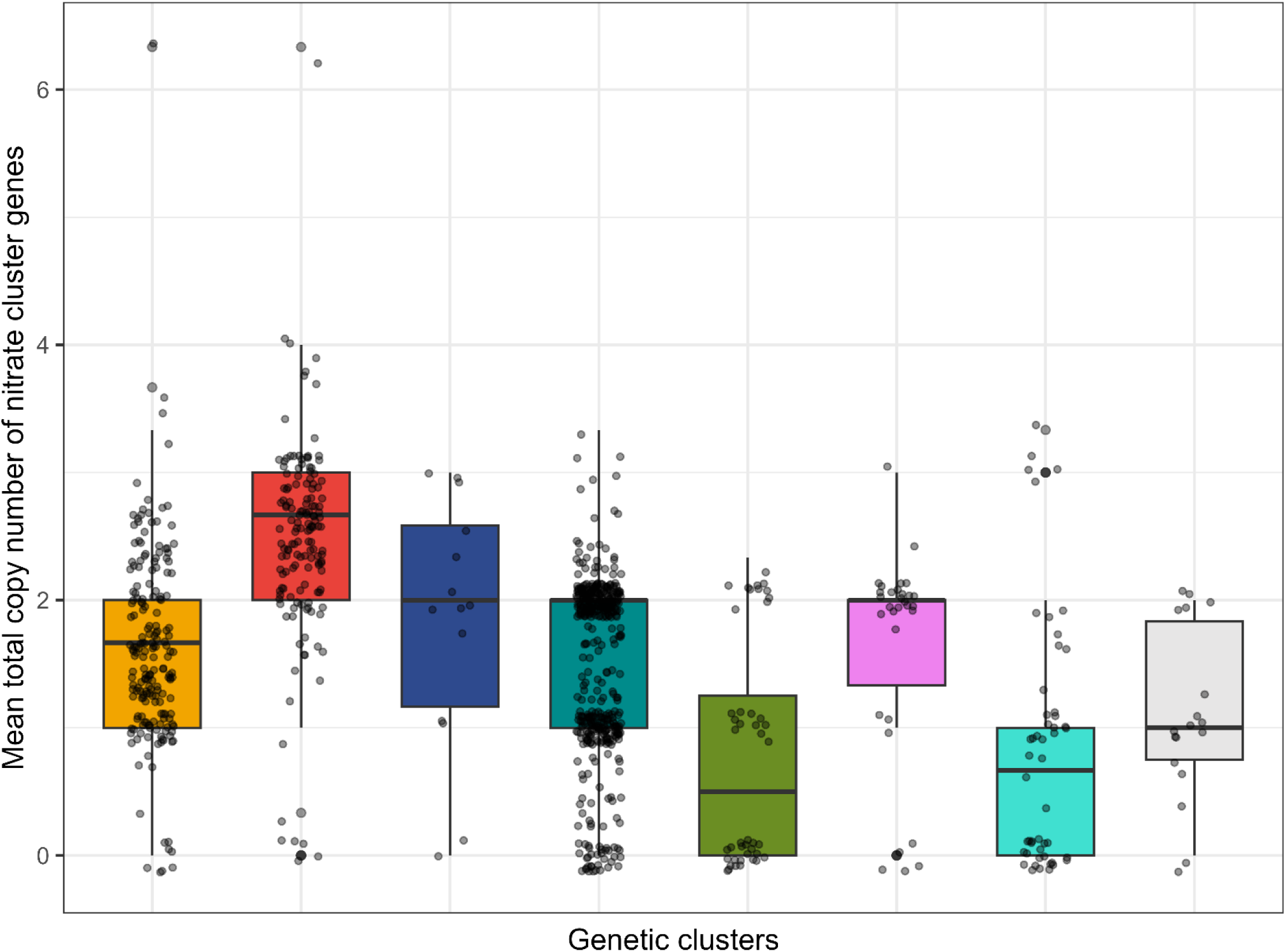
Mean total copy number of nitrate assimilation genes across genetic cluster. Distribution of the mean copy number of nitrate assimilation genes across genetic clusters, shown as boxplots. Each box represents the distribution of isolate-level mean copy numbers for a given genetic cluster. For allotriploid clusters, copy numbers from the primary and acquired genomes are summed.

**Supp.2.**
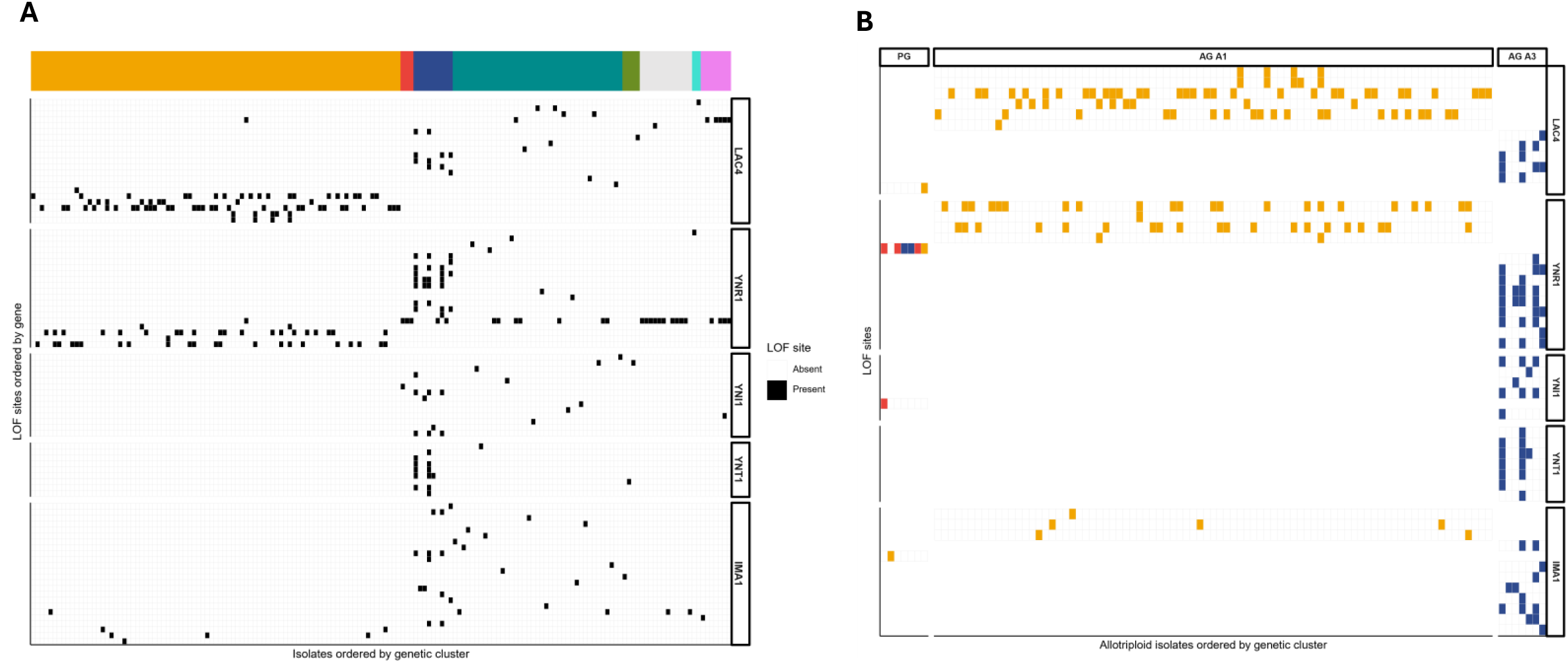
Distribution of loss-of-function (LOF) sites across isolates and subgenomes on chromosome IV. (A) Distribution of LOF sites across all isolates. Each column represents one isolate ordered by genetic cluster, and each row represents a distinct LOF site ordered by gene (LAC4, YNR1, YNI1, NT1, IMA1). Black tiles indicate the presence of a LOF variant in a given isolate at a given site, whereas white tiles indicate its absence. Only isolates carrying at least one LOF variant in the corresponding gene re represented. The colored annotation bar indicates the genetic cluster of each isolate. (B) Distribution of LOF sites in allotriploid isolates separated by subgenome. Columns represent isolates ordered by genetic cluster, and rows represent distinct LOF sites ordered by gene. Tiles indicate the presence of a LOF variant, colored according to the corresponding allotriploid genetic cluster (A1, A2, A3). Facets separate the primary genome (PG) from acquired genomes. Only isolates carrying at least one LOF ariant in the corresponding gene are represented.

**Table Supp.1.** Brettanomyces bruxellensis strains included in this study. List of the *B. bruxellensis* isolates used in this study, grouped according to their genetic cluster and ploidy level.

| *B. bruxellensis*<br>population | Ploidy | Number of strains | Strains |
| --- | --- | --- | --- |
| D1 | 2n | 44 | YJS5302, YJS5319, YJS5320, YJS5345, YJS5347,<br>YJS5357, YJS5373, YJS5385, YJS5392, YJS5422,<br>YJS5426, YJS5440, YJS5447, YJS5449, YJS5453,<br>YJS5458, YJS5461, YJS5479, YJS5485, YJS7800,<br>YJS7802, YJS7806, YJS7808, YJS7832, YJS7844,<br>YJS7848, YJS7849, YJS7856, YJS7860, YJS7862,<br>YJS7874, YJS7875, YJS7888, YJS7911, YJS7953,<br>YJS7954, YJS7955, YJS7967, YJS7973, YJS7990,<br>YJS8068, YJS8082, YJS8095, YJS8251 |
| Non-admixed D2 | 2n | 13 | YJS5310, YJS5340, YJS5344, YJS5363, YJS5368,<br>YJS5407, YJS5420, YJS7831, YJS7879, YJS7909,<br>YJS7910, YJS7923, YJS7947 |
| Admixed D2 | 2n | 16 | YJS5301, YJS5334, YJS5349, YJS5384, YJS5398,<br>YJS5402, YJS5406,<br>YJS5413, YJS5417, YJS5431, YJS7921, YJS7958,<br>YJS8013, YJS8070, YJS8090, YJS8285 |
| Admixed D1/D2 | 3n | 18 | YJS5382, YJS7801, YJS7804, YJS7807, YJS7809,<br>YJS7811, YJS7812, YJS7823, YJS7824, YJS7853,<br>YJS7891, YJS7894, YJS7989, YJS8000, YJS8045,<br>YJS8063, YJS8065, YJS8072 |
| A1 | 2n+1n | 20 | YJS5397, YJS5400, YJS5454, YJS7810, YJS7813,<br>YJS7816, YJS7817, YJS7818, YJS7858, YJS7868,<br>YJS7895, YJS7926, YJS7944, YJS7966, YJS7995,<br>YJS8010, YJS8048, YJS8081, YJS8089, YJS8229 |
| A2 | 2n+1n | 24 | YJS5408, YJS5434, YJS5445, YJS5459, YJS5469,<br>YJS5473, YJS5476, YJS5478, YJS5487, YJS7803,<br>YJS7827, YJS7838, YJS7881, YJS7886, YJS7900,<br>YJS7902, YJS7903, YJS7905, YJS7948, YJS7972,<br>YJS7980, YJS8091, YJS8092, YJS8757 |
| A3 | 2n+1n | 13 | YJS7805, YJS7819, YJS7820, YJS7821, YJS7828,<br>YJS7829, YJS7839, YJS7890, YJS7952, YJS8003,<br>YJS8004, YJS8014, YJS8028 |
| Admixed | 2n | 3 | YJS7884, YJS7885, YJS7889 |

**Table Supp.2.** Summary of loss-of-function (LOF) variants identified in nitrate cluster genes and flanking genes across genetic clusters and subgenomes. Counts of predicted LOF variants identified in nitrate assimilation genes (*YNR1, YNI1, YNT1*) and flanking genes (LAC4, IMA1) across genetic clusters and primary and acquired genomes in allotriploids. LOF variants include stop-gained, frameshift and start-loss mutations. For each genetic cluster, genome compartment and gene, the table reports the number of LOF events detected, the number of distinct LOF sites, and the number of isolates carrying at least one LOF variant.

| Genetic cluster | Subgenome | Gene | LOF type | Number of LOF events | Number of distinct LOF sites | Number of isolates carrying LOF |
| --- | --- | --- | --- | --- | --- | --- |
| A1 | AG | IMA1 | stop_gained | 6 | 3 | 6 |
| A1 | AG | LAC4 | frameshift | 38 | 4 | 33 |
| A1 | AG | LAC4 | stop_gained | 33 | 2 | 33 |
| A1 | AG | YNR1 | frameshift | 44 | 3 | 37 |
| A1 | AG | YNR1 | stop_gained | 1 | 1 | 1 |
| A1 | PG | IMA1 | stop_gained | 1 | 1 | 1 |
| A1 | PG | LAC4 | stop_gained | 1 | 1 | 1 |
| A1 | PG | YNR1 | start_lost | 1 | 1 | 1 |
| A2 | PG | YNI1 | stop_gained | 1 | 1 | 1 |
| A2 | PG | YNR1 | start_lost | 6 | 1 | 3 |
| A3 | AG | IMA1 | frameshift | 12 | 7 | 7 |
| A3 | AG | IMA1 | stop_gained | 1 | 1 | 1 |
| A3 | AG | LAC4 | frameshift | 9 | 4 | 4 |
| A3 | AG | LAC4 | stop_gained | 1 | 1 | 1 |
| A3 | AG | YNI1 | frameshift | 8 | 4 | 4 |
| A3 | AG | YNI1 | stop_gained | 1 | 1 | 1 |
| A3 | AG | YNR1 | frameshift | 23 | 8 | 5 |
| A3 | AG | YNR1 | stop_gained | 1 | 1 | 1 |
| A3 | AG | YNT1 | frameshift | 12 | 7 | 3 |
| A3 | PG | YNR1 | start_lost | 2 | 1 | 2 |
| Admixed | PG | IMA1 | stop_gained | 2 | 1 | 2 |
| Admixed | PG | LAC4 | stop_gained | 1 | 1 | 1 |
| Admixed | PG | YNR1 | start_lost | 12 | 1 | 10 |
| Admixed D1/D2 | PG | LAC4 | stop_gained | 1 | 1 | 1 |
| Admixed D1/D2 | PG | YNR1 | stop_gained | 1 | 1 | 1 |
| Admixed D2 | PG | IMA1 | stop_gained | 1 | 1 | 1 |
| Admixed D2 | PG | LAC4 | stop_gained | 5 | 1 | 5 |
| Admixed D2 | PG | YNI1 | stop_gained | 1 | 1 | 1 |
| Admixed D2 | PG | YNR1 | start_lost | 4 | 1 | 4 |
| D1 | PG | IMA1 | stop_gained | 12 | 11 | 12 |
| D1 | PG | LAC4 | stop_gained | 9 | 7 | 9 |
| D1 | PG | YNI1 | start_lost | 1 | 1 | 1 |
| D1 | PG | YNI1 | stop_gained | 6 | 6 | 6 |
| D1 | PG | YNR1 | start_lost | 8 | 1 | 7 |
| D1 | PG | YNR1 | stop_gained | 5 | 5 | 5 |
| D1 | PG | YNT1 | stop_gained | 1 | 1 | 1 |
| Non-admixed D2 | PG | IMA1 | stop_gained | 1 | 1 | 1 |
| Non-admixed D2 | PG | LAC4 | stop_gained | 1 | 1 | 1 |
| Non-admixed D2 | PG | YNI1 | stop_gained | 1 | 1 | 1 |
| Non-admixed D2 | PG | YNT1 | stop_gained | 1 | 1 | 1 |

